# Genetic analysis identifies molecular systems and biological pathways associated with household income

**DOI:** 10.1101/573691

**Authors:** W. David Hill, Neil M. Davies, Stuart J. Ritchie, Nathan G. Skene, Julien Bryois, Steven Bell, Emanuele Di Angelantonio, David J. Roberts, Shen Xueyi, Gail Davies, David C.M. Liewald, David J. Porteous, Caroline Hayward, Adam S. Butterworth, Andrew M. McIntosh, Catharine R. Gale, Ian J. Deary

## Abstract

Socio-economic position (SEP) is a multi-dimensional construct reflecting (and influencing) multiple socio-cultural, physical, and environmental factors. Previous genome-wide association studies (GWAS) using household income as a marker of SEP have shown that common genetic variants account for 11% of its variation. Here, in a sample of 286,301 participants from UK Biobank, we identified 30 independent genome-wide significant loci, 29 novel, that are associated with household income. Using a recently-developed method to meta-analyze data that leverages power from genetically-correlated traits, we identified an additional 120 income-associated loci. These loci showed clear evidence of functional enrichment, with transcriptional differences identified across multiple cortical tissues, in addition to links with GABAergic and serotonergic neurotransmission. We identified neurogenesis and the components of the synapse as candidate biological systems that are linked with income. By combining our GWAS on income with data from eQTL studies and chromatin interactions, 24 genes were prioritized for follow up, 18 of which were previously associated with cognitive ability. Using Mendelian Randomization, we identified cognitive ability as one of the causal, partly-heritable phenotypes that bridges the gap between molecular genetic inheritance and phenotypic consequence in terms of income differences. Significant differences between genetic correlations indicated that, the genetic variants associated with income are related to better mental health than those linked to educational attainment (another commonly-used marker of SEP). Finally, we were able to predict 2.5% of income differences using genetic data alone in an independent sample. These results are important for understanding the observed socioeconomic inequalities in Great Britain today.

---

People living in advantaged socio-economic backgrounds tend, on average, to live longer, and have better mental and physical health than those from more deprived environments.^1–3^ An understanding of the causes underlying the association between socioeconomic position (SEP) and health is likely to be helpful to minimize social disparities in health and wellbeing.^4^

The link between SEP and health is typically thought to be due to environmental factors including, but not limited to: access to resources, exposure to harmful or stressful environments, adverse health behaviors such as smoking, poor diet, and excessive alcohol consumption, and a lack of physical exercise.^5^ In addition, however, genetic factors have long been discussed as a partial explanation for the SEP-health association.^6^ It has recently been demonstrated that genome-wide association studies (GWASs) can capture shared genetic associations with both of these variables.^7^ Potential pleiotropic effects are highlighted in the observed genetic correlations between SEP variables such as completed years of education, household income, and social deprivation, and physical and mental health traits including longevity.^7, 8^

Loci associated with two SEP phenotypes, education and household income, have been identified via GWASs^7, 9–11^, but—consistent with other complex traits, such as height—these loci collectively account for only a small fraction of the total heritability of the traits in question. For education, 1,271 genomic loci were detected in the most recent meta-analysis of GWASs.^11^ For household income, analysis of a sample of 96,900 individuals from across the Great Britain found that additive genetic effects tagged by common SNPs accounted for approximately 11% (SE = 0.7%) of individual differences in household income.^7^ Two loci attained genome-wide significance in that study, but they collectively explained less than 0.005% of the total heritability.

Markers of SEP describe an individual’s social and economic position in the society in which they find themselves. As such, the link between genetic variation and SEP is not direct. The concept of mediated pleiotropy (sometimes referred to as vertical pleiotropy^12^) provides a possible explanation as to why an individual’s genotype is predictive of their income level. Mediated pleiotropy describes instances where one genetically-influenced phenotype is causally related to a second phenotype,^12, 13^ and therefore a genetic variant associated with the first phenotype (for instance a psychological trait such as intelligence, or conscientiousness, or a health trait such as disease resistance) might also be associated with a second, more biologically distal phenotype (in the case of the present study, income; Figure 1).

**Figure 1.**
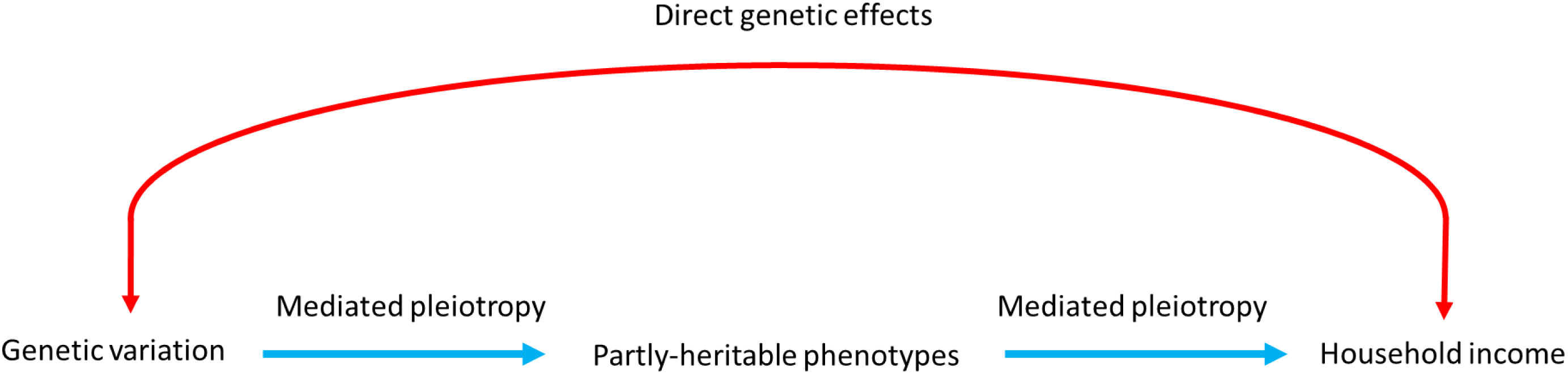
Illustrating the difference between a direct effect of genotype on income (shown in red), and the, more likely, notion that mediated pleiotropy (also termed vertical pleiotropy shown in blue) best explains the link between genetic variation and more biologically distal phenotypes. Mediated pleiotropy describes instances where genetic variation is linked to a phenotype (in this case household income) through genetic effects that act on another partly-heritable trait. These partly-heritable traits would also be associated with household income, and so the genetic effects that act on them would also be associated with household income. For simplicity, this schematic illustrates only a single link between genetic variation and household income. In reality there may be, and are likely to be, multiple links between genetic variation, and more biologically distal phenotypes such as household income.

Here, we use the UK Biobank dataset^14^ to examine genetic associations with household income (N=286,301) in a contemporary British sample. This study had four main aims. First, to identify genomic loci related to income and map these to genes, tissue types, and biological processes that may help to elucidate potential mechanisms linking income to inequalities in health in the UK today. Second, since differences in income have been linked to differences in cognitive ability (also called intelligence), both using phenotypic data^15, 16^ and molecular genetic analyses,^7^ we use two-sample Mendelian Randomization (MR) to explore the causal links between intelligence and income. Because genetic variants are randomly “assigned” to children at conception, under a number of assumptions^17, 18^ MR can be seen as analogous to a randomised control trial where, in the present context, intelligence (operationalized using SNPs that attained genome-wide significance in a GWAS of intelligence) is randomly assigned to a participant at conception. This random allocation of intelligence at conception can be used to indicate the causal relationship between intelligence and income. Third, we use Multi-trait-based conditional & joint analysis (mtCOJO) to explore whether the genetic effects linked to intelligence explain the genetic association of income with mental health, metabolic, health and wellbeing, anthropometric, and reproductive traits, as well as other indicators of SEP. Fourth, we use polygenic risk scores^19^ derived from our discovery analyses to predict income using only DNA in an independent study.

Owing to the substantial genetic correlation between income and education (r_g_ = 0.73) that was found in a previous study,^8^ we use Multi-Trait Analysis of Genome-wide association studies (MTAG) to facilitate the discovery of additional loci and improve genetic prediction.^20^ MTAG is used to conduct a meta-analysis using summary statistics from genetically correlated traits, adding power in order to identify associations specific to one phenotype under investigation. In the current study, we use MTAG to meta-analyze a GWAS on household income performed in UK Biobank with a GWAS on years of education from a previous GWAS on Education.^10^ By combining these data, we increased the power of our GWAS of household income, attaining an effective sample size of 505,541 participants.

## Method

### Participants

The primary sample used involved participants from UK Biobank, an open-access resource established to examine the determinants of disease in middle-aged and older adults living in the United Kingdom.^21^ Recruitment to UK Biobank occurred between 2006 and 2010, targeting community-dwelling individuals from both urban and rural environments across a broad range of socio-economic circumstances. A total of 502,655 participants were assessed at baseline on a range of cognitive and other psychological measures, physical and mental health, and their socioeconomic position. They donated a number of biological samples, including DNA for genotyping. In order to reduce the effects of population stratification, only participants from a single ancestry group, those of White British ancestry, were included in the analysis. High quality genotyping was performed on 332,050 participants.

### Phenotype description

A total of 332,050 participants had genotype data and data on their level of household income. Self-reported household income was collected using a 5 point scale corresponding to the total household income before tax, 1 being less than £18,000, 2 being £18,000 - £29,999, 3 being £30,000 - £51,999, 4 being £52,000 – £100,000, and 5 being greater than £100,000. Participants were removed from the analysis if they answered “do not know” (n = 12,721), or “prefer not to answer” (n = 31,947). This left a total number of 286,301 participants (138,425 male) aged 39-73 years (mean = 56.5, SD = 8.0 years) with genotype data who had reported, between 1 and 5, their level of household income.

### UK Biobank genotyping

Full details of the UK Biobank genotyping procedure have been made available.^22^ In brief, two custom genotyping arrays were used to genotype 49,950 participants (UK BiLEVE Axiom Array) and 438,427 participants (UK Biobank Axiom Array).^22, 23^ Genotype data on 805,426 markers were available for 488,377 of the individuals in UK Biobank. Imputation to the Haplotype Reference Consortium (HRC) reference panel lead to 39,131,578 autosomal SNPs being available for 487,442 participants.^22^ Allele frequency checks^24^ against the HRC^25^ and 1000G^26^ site lists were performed, and variants with minor allele frequencies (MAF) differing more than +/− 0.2 from the reference sets were removed.

Additional quality control steps were conducted and described previously.^8, 27^ These included the removal of those with non-British ancestry based on self-report and a principal components analysis, as well as those with extreme scores based on heterozygosity and missingness. Individuals with neither XX or XY chromosomes, along with those individuals whose reported sex was inconsistent with genetically inferred sex, were also removed, as were those individuals with >10 putative third degree relatives from the kinship table. Following these exclusions, a sample of 408,095 individuals remained. Using GCTA-GREML on 131,790 reportedly-related participants,^28^ related individuals were removed based on a genetic relationship threshold of 0.025. Following this quality control, household income data, and genetic data, were available on 286,301 participants. Following association analysis, SNPs with a minor allele frequency < 0.0005, and an imputation quality score < 0.1 were removed. Finally, only bi-allelic SNPs were retained, resulting in 18,485,882 autosomal SNPs.

### Statistical analysis

A flow chart summarizing all statistical analyses conducted is displayed in Figure 2. These include:

**Figure 2.**
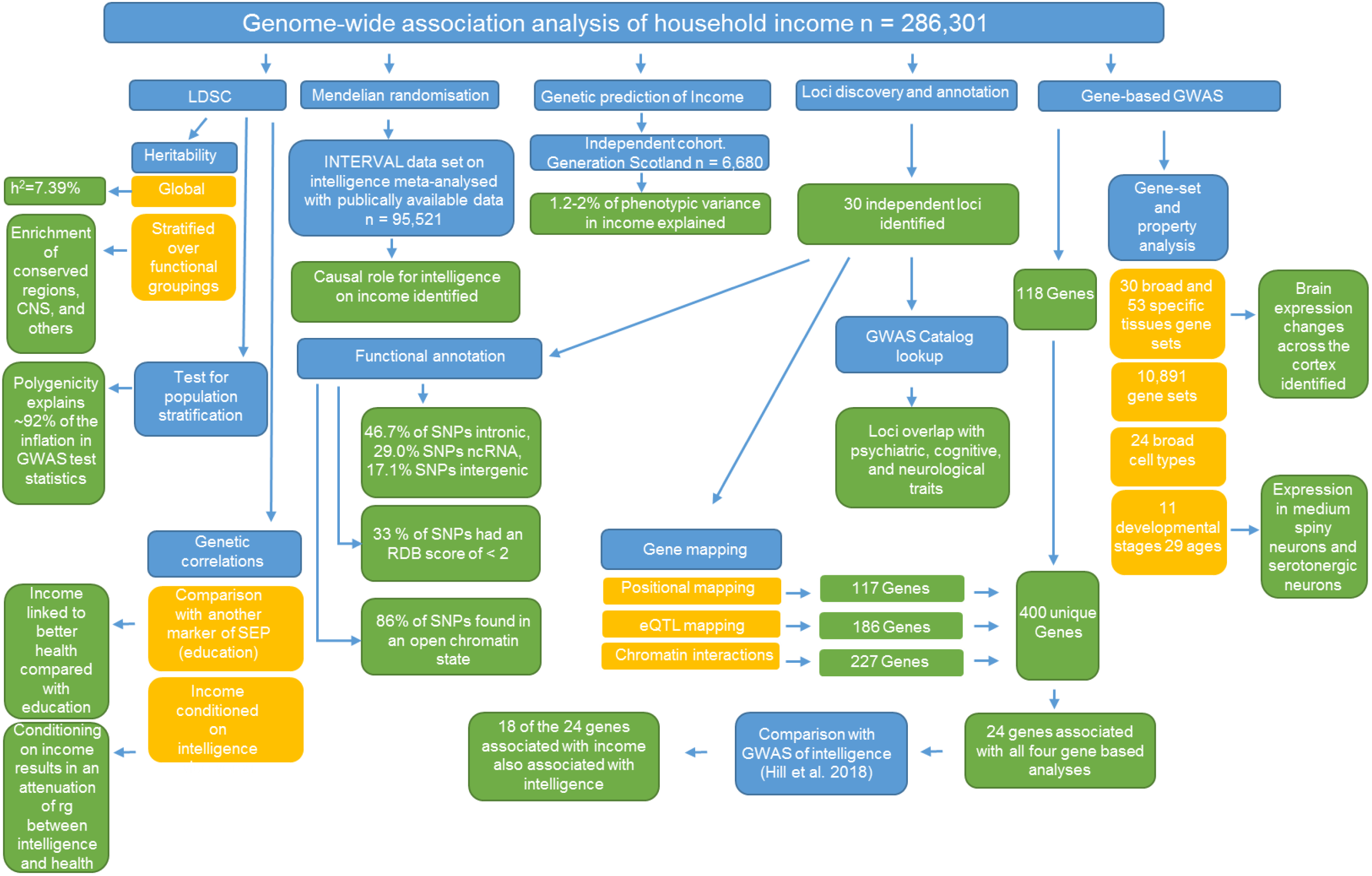
Flow chart for the statistical analysis carried out using the GWAS data on household income in 286,301 White British participants in UK Biobank. Blue indicates a type of analysis conducted (i.e. LDSC to derive a heritability estimate) and gold indicates a subtype of this type of analysis (i.e. global heritability or the heritability of a stratified subset of the SNPs). Green indicates the result of an analysis (i.e. the global heritability was 7.39%).

### Genome-wide association analysis (GWAS) in the UK Biobank sample

The level of household income as measured on the 5 point scale was subjected to a regression using income as the outcome as has been conducted previously,^7^ and 40 genetic principal components (to control for population stratification), genotyping array, batch, age, and sex as predictors. The residuals from this model were then used in a GWAS assuming an additive genetic model as implemented in BGENIE.^22^

### Functional annotation and loci discovery

Genomic risk loci were derived using the summary data from the data set of household income derived in UK Biobank, using FUnctional Mapping and Annotation of genetic associations (FUMA)^29^. FUMA was first used to identify independent significant SNPs using the *SNP2GENE* function. SNPs with a P-value of ≤ 5 ×10^−8^ and independent of other genome wide significant SNPs at r^2^ 0.6 were first identified. Using these independent significant SNPs, additional candidate SNPs, used in subsequent annotations, were defined as all SNPs that had a MAF of 0.001 and were in LD of ≥ r^2^ 0.6 with at least one of the independent significant SNPs. These candidate SNPs included those from the HRC panel and may not have been included in the GWASs performed on household income. Lead SNPs were also identified using the independent significant SNPs. Lead SNPs were defined as SNPs that were independent from each other at r^2^ 0.1. Next, genomic risk loci that were 250kb or closer were merged into a single locus.

The lead SNPs and those in LD with the lead SNPs were then mapped to genes based on their functional consequences, as described using ANNOVAR^30^ and the Ensemble genes build 85. Intergenic SNPs were annotated as the two closest flanking genes which can result in them being assigned to multiple genes.

### Gene-mapping

Three strategies were used to link the income-associated independent genomic loci to genes. First, positional mapping^31^ was used to map SNPs to genes based on physical distance. SNPs were mapped to genes if they were within a 10kb from a known protein gene found in the human reference assembly (hg19).

Second, expression quantitative trait loci (eQTL) mapping was carried out by mapping SNPs to genes if allelic variation at the SNP is associated with expression levels of a gene. For eQTL mapping, information on 45 tissue types from three data bases (GTEx v7, Blood eQTL browser, BIOS QTL browser) based on cis-QTLs was used and SNPs were mapped to genes up to 1Mb away. A false discovery rate (FDR) of 0.05 was used as a cut off to define significant eQTL associations.

Finally, chromatin interaction mapping was carried out to map SNPs to genes when there is a three-dimensional DNA-DNA interaction between the SNP and gene. No distance boundary was applied as chromatin interactions can be long-ranging and span multiple genes. Hi-C data of 14 tissue types was used for chromatin interaction mapping.^32^ In order to reduce the total number of genes mapped using chromatin interactions and to increase the likelihood that those mapped are biologically relevant, an additional filter was added. We only retained interaction mapped genes if one region involved with the interaction overlapped with a predicted enhancer region in any of the 111 tissue/cell types found in the Roadmap Epigenomics Project,^33^ and the other region was located in a gene promoter region (i.e., 250bp upstream and 500bp downstream of the transcription start site and also predicted to be a promoter region by the Roadmap Epigenomics Project ^33^). An FDR of 1×10^−5^ was used to define a significant interaction.

### Gene-based GWAS

Gene-based analyses have been shown to increase the power to detect association due to the multiple testing burden being reduced, in addition to the effects of multiple SNPs being combined.^34^ Gene-based GWAS was conducted using MAGMA^35^. All SNPs located within protein coding genes were used to derive a P-value describing the association found with household income. The NCBI build 37 was used to determine the location and boundaries of 18,782 autosomal genes and linkage disequilibrium within and between genes was gauged using the HRC panel. In order to control for multiple testing, a Bonferroni correction was applied using each gene as an independent statistical unit (0.05 / 18,782 =2.66 ×10^−6^). The gene-based statistics derived using MAGMA were then used to conduct the gene-set analysis, the gene-property analyses, and the cell type enrichment analysis.

### Gene-set analysis

In order to understand the biological systems vulnerable to perturbation by common genetic variation, a competitive gene-set analysis was performed. Competitive testing, conducted in MAGMA,^35^ examines if genes within the gene-set are more strongly associated with the trait of interest than other genes, and differs from self-contained testing by controlling for type 1 error rate as well as being able examine the biological relevance of the gene-set under investigation.^36^

A total of 10,891 gene-sets (sourced from Gene Ontology,^37^ Reactome,^38^ and, MSigDB^39^) were examined for enrichment of household income. A Bonferroni correction was applied to control for the multiple tests performed on the 10,891 gene-sets available for analysis.

### Gene-property analysis

In order to identify the relative importance of particular tissue types which may indicate the intermediary biological phenotypes that might act between genetic variation and SEP outcomes, a gene property analysis was conducted using MAGMA. The goal of this analysis was to determine if, in 30 broad tissue types, and 53 specific tissues, tissue specific differential expression levels were predictive of the association of a gene with household income. Tissue types were taken from the GTEx v7 RNA-seq database^40^ with expression values being log2 transformed with a pseudocount of 1 after Winsorising at 50 with the average expression value being taken from each tissue. Multiple testing was controlled for using Bonferroni correction. An additional gene property analysis was conducted to determine if transcription in the brain at any one of 11 developmental stages,^41^ or across 29 different specific ages,^41^ was associated with a gene’s link to household income. A Bonferroni correction was used to control for 11 and 29 tests separately.

### Cell type enrichment

As previous studies had indicated the importance of cortical tissues to differences in SEP,^7, 10^ a gene property analysis was also conducted to examine a broad array of brain specific cell types. Enrichment of heritability was tested against 173 types of brain cells (24 broad categories of cell types), which were calculated following the method described in Skene et al^42^. Briefly, brain cell-type expression data were drawn from single-cell RNA sequencing data from mouse brains. For each gene, a specificity value for each cell-type was calculated by dividing the mean Unique Molecular Identifier (UMI) counts for the given cell type by the summed mean UMI counts across all cell types. MAGMA^35^ was used to calculate cell type enrichments where specificity values were then divided into 40 equal sized bins for each cell type for the MAGMA analysis. A linear model was fitted over the 40 specificity bins (with the least specific bin indexed as 1 and the most specific as 40). This was done by passing the bin values for each gene using the ‘*--gene-covar onesided*’ argument.

### Univariate linkage disequilibrium score

Univariate LDSC regression was performed on the summary statistics from the GWAS on household income in order to quantify the degree to which population stratification may have influenced these results.

For the GWAS on household income, an LD regression was carried out by regressing the GWA test statistics (χ^2^) from each GWAS onto the LD score (the sum of squared correlations between the minor allele frequency count of a SNP with the minor allele frequency count of every other SNP) of each SNP. This regression allows for the estimation of heritability from the slope, and a means to detect residual confounders using the intercept.

LD scores and weights were downloaded from (http://www.broadinstitute.org/~bulik/eur_ldscores/) for use with European populations. A minor allele frequency cut-off of > 0.1 and an imputation quality score of > 0.9 were applied to the GWAS summary statistics. Following this, SNPs were retained if they were found in HapMap 3 with MAF > 0.05 in the 1000 Genomes EUR reference sample. Following this indels and structural variants were removed along with strand ambiguous variants. SNPs whose alleles did not match those in the 1000 Genomes were also removed. As the presence of outliers can increase the standard error in LDSC score regression and so SNPs where χ^2^ > 80 were also removed.

### Partitioned heritability

Partitioned heritability was performed using stratified linkage disequilibrium score (LDSC) regression.^43^ Partitioned heritability analysis aims to determine if SNPs that account for variance in income cluster in functional regions of the genome. Full details of how this method works can be found in Finucane et al.^43^ Firstly, heritability for each of the functional groups is derived. Secondly, this heritability estimate is used to compute an enrichment metric that is defined as the proportion (Pr) of heritability captured by the functional annotation, over the proportion of SNPs contained within the functional annotation (Pr(h^2^)/Pr(SNPs)). This ratio describes if a functional annotation contains a greater or lesser proportion of the heritability than would be predicted by chance, with chance being defined as the proportion of heritability explained being equal to the proportion of SNPs within the functional annotation (Pr(h^2^)/Pr(SNPs) = 1).

Stratified LD Scores were calculated from the European-ancestry samples in the 1000 Genomes project (1000G) and only included the HapMap 3 SNPs with a minor allele frequency (MAF) of >0.05. The model was constructed using 52 overlapping, functional categories. Correction for multiple testing was performed using a Bonferroni test on the 52 functional categories (α = 0.00096).

### Mendelian Randomization

The causal effects of intelligence (termed the “exposure” in an MR analysis) on income (termed the “outcome” in an MR analysis) were investigated using univariate Mendelian Randomization (MR) analysis. Here, the total causal effect of intelligence on income was examined by combining summary GWAS test statistics for intelligence and for income using an inverse-variance-weighted (IVW) regression model.^44^ This is equivalent to a weighted regression of the SNP-outcome coefficients on the SNP-exposure coefficients, with the intercept constrained to zero (i.e. assuming no or balanced horizontal pleiotropy).

The results of the IVW regression model were compared with the results obtained using MR-Egger regression,^45^ due to the use of multiple alleles in MR analyses increasing the potential for pleiotropic effects caused by the aggregation of invalid genetic instruments.^46^ By not constraining the intercept to zero (as done using inverse variance weighted regression) MR-Egger relaxes the assumption that the effects of genetic variants on the outcome act solely through the exposure (in this case intelligence). The intercept parameter of the MR-Egger regression indicates the average directional pleiotropic effects of the SNPs on the outcome. As such, the direct pleiotropic effect that the SNPs have on the outcome, independent of the exposure, can be quantified, where a non-zero intercept provides evidence for bias due to directional pleiotropy and a violation of the MR IVW estimator assumptions. Of note is that the MR-Egger regression estimates only remain consistent if the magnitude of the gene-exposure associations, across all variants, are independent of their pleiotropic effects (i.e. the InSIDE assumption holds).^45^ In addition, power is almost always lower for MR-Egger and it requires variation in the effects of the SNPs on the exposure (i.e. if all SNPs have similar effects on the exposure, then MR-Egger will have very low power).

For use with Mendelian Randomization, two-independent groups (n = 95,521 for intelligence and n = 271,732 for income) were created whereby the GWAS on income was re-run using only those participants whose data were not included in the interim release of the UK Biobank genotype data. A GWAS data set on intelligence was created by meta-analysing publicly-available data on intelligence with a new GWAS (conducted for this study) on intelligence using data from the INTERVAL BioResource^47, 48^ (**Supplementary data**) where 19 SNPs were identified as being genome-wide significant and independent. These 19 SNPs were used as instrumental variables for intelligence in the MR analysis.

### Genetic correlations

Genetic correlations were derived using bi-variate LDSC regression. The data processing pipeline used by Bulik-Sullivan et al.^49^ was used and 26 GWAS data sets on health, anthropometric, psychiatric, cognitive, and metabolic traits were selected (**Supplementary Table 1**). There were three objectives to our analysis examining genetic correlations using household income. First, we sought to replicate the results of Hill et al. (2016)^7^ who found genetic correlations between household income and other variables in a smaller data subset from the UK Biobank sample used here. Second, SEP is multi-dimensional in nature: it is composed of multiple measures, each of which are correlated imperfectly with the others. Because of this, different measures of SES may have genetic variance that is both unique to them, and differentiates them from the others in the way it associates with health. To examine this, we compare how genetic correlations with household income and 26 health, anthropometric, psychiatric, cognitive, and metabolic traits differed compared to the genetic correlations derived using a different, individual-level measure of SEP, i.e. educational attainment as measured by the number of years one has spent in education.^11^ Third, Hill et al. (2016) also found that the phenotypes with the strongest genetic correlations with income are those that are “cognitive” (verbal numerical reasoning, childhood IQ, and years of education) in nature.^7^ The magnitude of these genetic correlations might indicate the phenotypes that occur as potential mediators between molecular genetic inheritance and household income.

In addition, intelligence is known to be genetically correlated with many physical and mental health traits.^50–52^ The role that intelligence might play in accounting for some of the genetic links between household income and 26 health and wellbeing, anthropometric, mental health, and metabolic traits was examined using genetic correlations. Here, the GWAS of income was conditioned on a GWAS on intelligence using Multi-trait-based conditional & joint analysis (mtCOJO). mtCOJO is used to perform conditional GWAS where by the genetic effects from one GWAS are controlled for in another GWAS. Importantly, the mtCOJO method avoids well-known issues of collider bias that can occur by including heritable covariates.^53^ In the current study, the GWAS on income was conditioned on a GWAS on intelligence (and the intelligence GWAS was conditioned on the income GWAS) before the genetic correlations between income (and intelligence) and 26 variables mentioned above were re-ran.

### Genetic prediction

Using the summary statistics from our GWAS of household income polygenic risk scores (PGRS) were derived using PRSice-2^54^ and the Generation Scotland: Scottish Family Health Study (GS:SFHS) cohort. The recruitment protocol and sample characteristics of GS:SFHS are described in full elsewhere.^55, 56^ In brief, 23,690 participants were recruited through their GP from across Scotland. Participants were all aged 18 and over and were not ascertained based on the presence of any specific disease. Following the removal of individuals who preferred not to answer, income was assessed in GS:SFHS by 5 point scale (1 less than £10,000, 2 between £10,000 and £30,000, 3 between £30,000 and £50,000, 4 between £50,000 and £70,000, 5 more than £70,000). Individuals who preferred not to answer were excluded from the analysis. Individuals who had taken part in UK Biobank were also removed from the GS:SFHS data set (n = 174). SNPs were included in the data if they had a MAF of ≥ 0.01 and Hardy-Weinberg P-value of 0.000001. Finally, one from every pair of related individuals were removed from the data set by creating a genetic relationship matrix using GCTA^57^ and removing individuals who are related at ≥ 0.025. This yielded a final sample size of 6,680 participants who had genotype data and income data.

The participant’s level of income was then used as a predictor in a regression analysis with age, sex, and 20 principal components included to control for population stratification. The standardized residuals from this model were then used as each participant’s income phenotype. PGRS were created using the income phenotype derived using UK Biobank.

In each instance PRSice-2 was used to create five PGRS corresponding to one of five P-value cut-offs (P ≤ 0.01, P ≤ 0.05, P ≤ 0.1, P ≤ 0.5, P ≤ 1) applied to the association statistics from the summary data. The polygenic risk scores for each threshold were then standardized and used in a regression model to predict the income phenotype in GS:SFHS.

### Multi-Trait Analysis of GWAS (MTAG)

MTAG^20^ can be used to meta-analyze genetically correlated traits in order to increase power to detect loci in any one of the traits. Only summary data are required in order to carry out MTAG and bivariate LD score regression is carried out as part of an MTAG analysis to account for (possibly unknown) sample overlap between the GWAS data sets.^20^ The goal of this analysis was to increase the power to detect loci associated with income, and so our income GWAS was meta-analysed with the GWAS on years of education by Okbay et al.^58^ using MTAG. Both the Okbay data set and the income data set from UK Biobank had a similar level of power (Okbay mean χ^2^ = 1.65, UK Biobank income mean χ^2^ = 1.45) and they showed a genetic correlation of r_g_ = 0.77 (SE = 0.02), confirming that both income and education, as measured using these data sets, have a highly similar genetic etiology.

Functional annotation and loci discovery, gene-mapping, gene-based GWAS, gene-set and gene-property analysis, were also performed using the MTAG derived data set on income. In addition, following the removal of UK Based cohorts from the educational attainment summary statistics, genetic prediction was performed using the MTAG derived income phenotype and the GS:SFHS as described above.

## Results

### SNP-based analysis of income

For household income, 3,712 SNPs attained genome-wide significance (*P* < 5×10^−8^), across 30 independent loci (Figure 3A & **Supplementary Table 2**) which contained 68 independent significant SNPs and 31 lead SNPs. A total of 29 of these 30 loci were not reported in the previous UK Biobank analysis of income and should therefore be considered novel^7^ (**Supplementary Table 3**). The 30 loci predominantly contained SNPs found within intronic regions (47%) as well as non-coding RNA introns (29%). A total of 17% of the SNPs within the independent loci were found in intergenic regions, and only 1.2% were found in exons (Figure 3B). Many of the loci contained SNPs showing evidence of influencing gene regulation with 33% having a Regulome-DB score of <2 (Figure 3C) and 86% having evidence of being in an open chromatin state (indicated by a minimum chromatin state of <8, in Figure 3D).

**Figure 3A.**
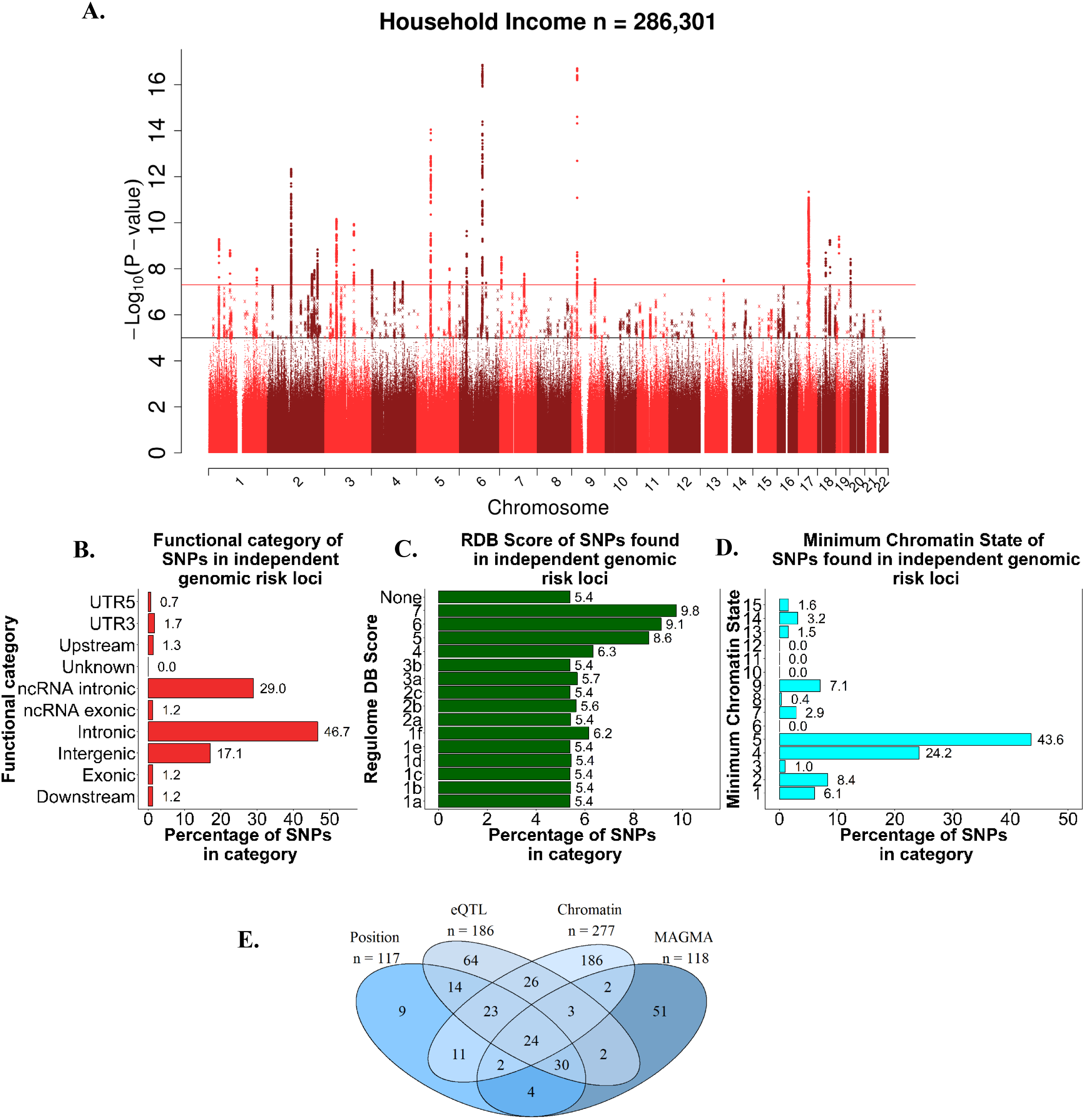
Manhattan plot for income; negative log10 transformed P-values for each SNP are plotted against chromosomal location. The red line indicates genome-wide significance and the black line indicates suggestive associations (1 × 10^−5^). **Figure 3B**. Functional annotation carried out on the independent genomic loci identified. The percentage of SNPs found in each of the nine functional categories is listed. **Figure 3C**. The percentage of SNPs from the independent genomic loci that fell into each of the Regulome DB scores categories. A lower score indicates greater evidence for that SNPs involvement in gene regulation. **Figure 3D**. The percentage of SNPs within the independent genomic loci plotted against the minimum chromatic state for 127 tissue/cell types. **Figure 3E**. Venn diagram illustrating the overlap of the genes implicated using positional mapping, eQTL mapping, chromatin interaction mapping, that was conducted on the independent significant loci identified in the SNP-based GWAS. Also shown is how these implicated genes overlap with those identified using the gene-based statistics derived using MAGMA.

Using GWAS catalogue, six of these 30 loci associated with household income have previously been associated with intelligence. Eleven have been previously linked to education.^10^ Some have been linked with psychiatric diseases such as schizophrenia (1 locus),^59^ and bipolar disorder (2 loci).^60^ Some have been linked with the personality trait neuroticism (4 loci).^27^ Some have been linked with neurological variables (corticobasal degeneration, 1 locus; subcortical brain volumes, 1 locus), and Parkinson’s disease, (1 locus), reproductive traits (age of first birth, 2 loci), and other disease states (Crohn’s disease, 1 locus; inflammatory bowel disease, 2 loci; coronary artery disease, 2 loci). (**Supplementary Table 4**).

Linkage disequilibrium score (LDSC) regression was carried out on the GWAS summary data for household income. The mean χ^2^ statistic was 1.45 and the intercept of the LDSC regression was 1.04. These statistics indicate that around 92% of the inflation in the GWAS test statistics was due to a polygenic signal rather than residual stratification or confounding. It is important to note that, whereas LDSC regression can capture the effects of population stratification and cryptic relatedness, it is not expected to capture the genetic associations with parental income that might influence the environment a child was reared in. Should these indirect genetic effects^61^ play role in income differences they will remain in the GWAS estimates. The LDSC regression estimate of the heritability of household income was 7.39% (SE=0.33%). Whereas this is lower than previous estimates,^7^ it is known that, relative to GCTA-GREML, LDSC regression can produce lower estimates.^62^

### Gene prioritization of SNP based analysis of income

Three methods of mapping allelic variation to genes were used to better understand the functional consequences of the 30 independent loci linked to household income (positional mapping, eQTL analysis, and chromatin mapping). Using positional mapping, SNPs from the GWAS were aligned to 117 genes. eQTL mapping was used to match cis-eQTL SNPs to 186 genes, and chromatin interaction mapping linked the SNPs to 277 genes (Figure 3E & **Supplementary Table 5**). These mapping strategies identified a total of 400 unique genes, of which 133 (Figure 3E cells 14+23+26+3+24+11+2+30) were implicated by at least two mapping strategies and 47 (Figure 3E cells 23+24) were implicated by all three. Of the 133 implicated by two mapping strategies, two showed evidence of a chromatin interactions with two independent genomic risk loci (**Supplementary Table 6**). Both *HOXB2* and *HOXB7* showed interactions with loci 24 and loci 25. *HOXB2* showed interactions in mesendoderm (an embryonic tissue layer) tissue and IMR90 (fetal lung fibroblasts) tissue, whereas *HOXB7* showed associations in the tissues of hESC (human embryonic stem cell), Mesenchymal (multipotent stromal cells which differentiate into a variety of different cell types) Stem Cell, IMR90, Left Ventricle, GM12878, and Trophoblast-like Cells.

### Gene-based analysis of income

In a gene-based GWAS using MAGMA,^63^ 118 genes were associated with income (P < 2.662×10^−6^) (**Supplementary Table 7** & Figure 4A). These 118 associated genes overlapped with 24 of those implicated using positional, eQTL, and chromatin interaction modelling (Figure 3E). Of the genes implicated by each of the three methods and the gene-based-GWAS, *BSN* was of particular note due to its being expressed primarily in the neurons of the brain and its role in the scaffolding protein involved in the organization of the presynaptic cytoskeleton. *MAPT* was also noted due to MAPT transcripts being differentially expressed in the nervous system dependent on the level of maturation and type of neuron in which it is found. In addition, mutations in the *MAPT* gene have also been linked to Alzheimer’s disease, frontotemporal dementia, cortico-basal degeneration, and progressive supranuclear palsy, all diseases with the common theme of neurodegeneration. Also found in this overlap was the gene *CHST10*. The protein encoded by this gene is a sulfotransferase that acts on HNK-1 which is involved in neurodevelopment and synaptic plasticity.

**Figure 4A.**
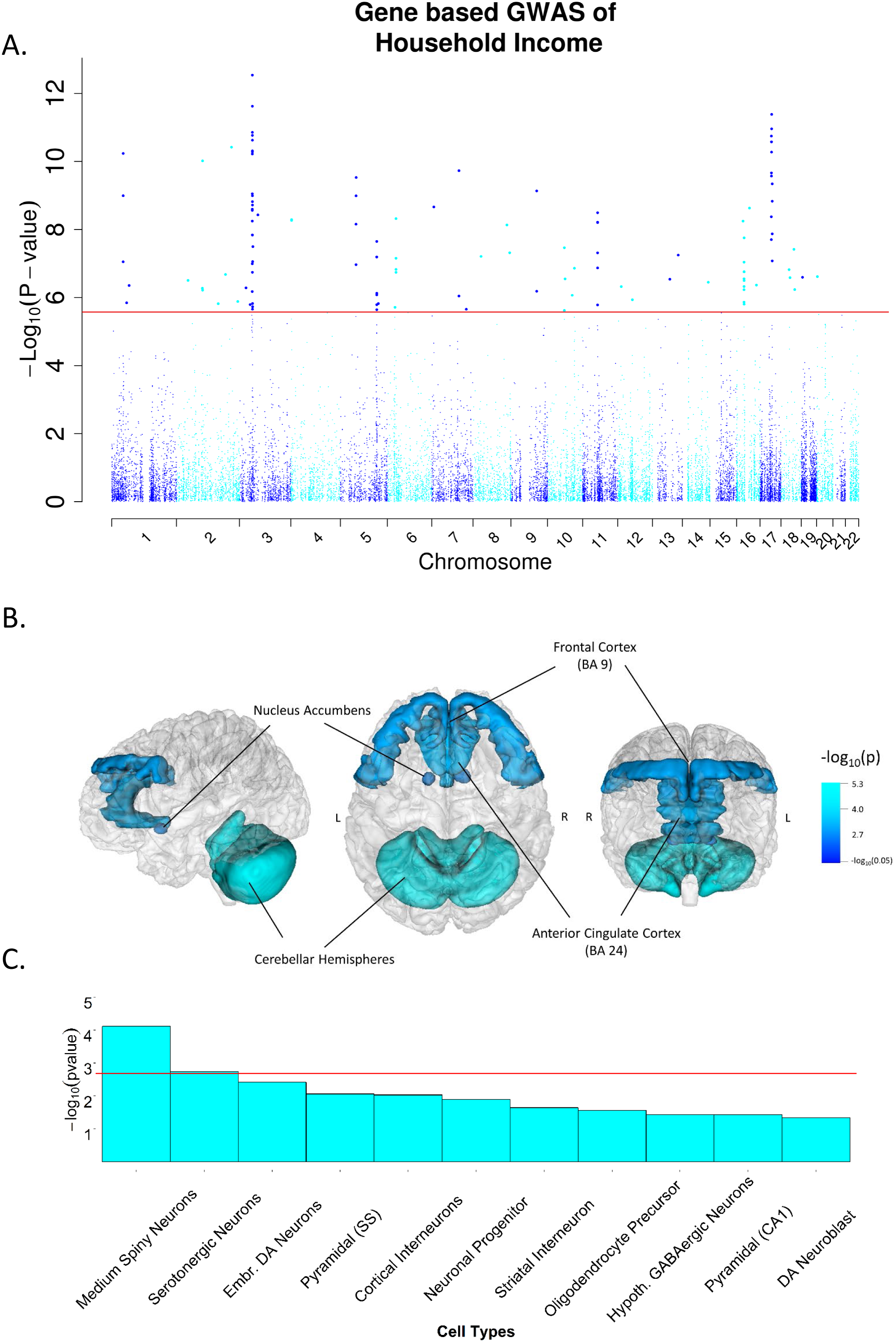
A Manhattan plot of income using the gene-based statistics derived using MAGMA; negative log10 transformed P-values for each gene are plotted against chromosomal location. The red line indicates genome-wide significance. **Figure 4B**. The results of a gene-property analysis linking transcription differences in the brain with income differences. Significant links between expression differences in cerebellar hemisphere, frontal cortex (Brodmann area 9), the nucleus accumbens and the anterior cingulate cortex (Brodmann area 9 24) are illustrated. Dark blue indicates low –log_10_ P-values (a lower level of association) describing the link between gene expression and household income and light blue indicates high –log_10_ P-values (a higher level of association) describing the same relationship. The full results found in **Supplementary Table 9**) with the gene based statistics produced using MAGMA. **Figure 4C**. Shows the results of a cell type specific gene-property analysis where the relationship between the gene-based statistics from MAGMA and the degree to which gene expression was specific to the annotations was examined. A Bonferroni correction was applied to control for the 24 tests conducted. The red line indicates statistical significance indicating that expression that is specific to the annotation is associated with the gene-based statistics for income.

These 24 genes were then examined to determine if gene-based statistics had implicated them in intelligence due to the previously-reported, strong genetic correlations between income and intelligence.^7^ We found that 18 were associated (P<2.75×10^−6^) with intelligence in a previous gene-based analysis.^8^ This indicates that the genes with the most biological relevance to income were also linked to intelligence, again suggestive of the role that intelligence plays in SEP differences. Four of these, *BSN*, *RBM5*, *KANSL1*, *AFF3*, and *CAMKV*, are each highly intolerant to loss of function mutations as indicated by their probability of loss-of-function score being > 0.9. Of the 6 that were not genome-wide significant, the levels of association ranged from P=5.75 ×10^−6^ to 0.001.

### Gene-set and gene-property analysis of income

Gene-set analysis did not find evidence that any of the gene-sets included here were enriched for differences in household income (**Supplementary Table 8**). However, a gene-property analysis showed that genes that were more associated with household income in the MAGMA analysis were also more highly expressed in the brain (P=1.31×10^−5^) and the testis (P=1.31×10^−5^) than genes that were less associated with income (**Supplementary Table 9**). This relationship was found across tissues in the cerebellum (P=5.61×10^−6^), the cerebellar hemisphere (P=5.99×10^−6^), the frontal cortex BA9 (P=9.68×10^−5^), the cortex (P=1.05×10^−4^), the nucleus accumbens basal ganglia (P=2.93×10^−4^), and the anterior cingulate cortex BA24 (P=6.81×10^−4^) (**Supplementary Table 10** & Figure 4B).

Cell-type analysis conducted on household income indicated that, of the 24 cell types examined, two were statistically significant after controlling for 24 tests. The significant cell types include medium spiny neurons P=7.67×10^−5^, and serotonergic neurons P=0.002 (**Supplementary Table 11** & Figure 4C). Finally, gene-property analysis found little evidence that genes linked to household income were transcribed in the brain at any one of 11 developmental stages,^41^ or across 29 different specific ages^41^ (**Supplementary Table 12** & **Supplementary Table 13**).

### Partitioned heritability

The results of the partitioned heritability analysis provide a complementary means to examine the biological relevance of the signal captured by the GWAS of income. The partitioned heritability analysis examines all SNPs in the GWAS, without utilizing a P-value cut off. In this way it differs from the functional annotation of the SNP-based analysis, the gene prioritization, and the gene-based GWAS. The partitioned heritability analysis describes whether or not the SNPs that capture the greatest proportion of the heritability of income, also cluster in regions of the genome that are united by a shared biological theme. We find that, consistent with the notion that intelligence and income are genetically linked,^64^ the regions of the genome that have undergone purifying selection are those that harbor the greatest proportion of heritability for income. These regions contained only 2.6% of the SNPs but collectively accounted for 53.9% of the total heritability as derived using LDSC regression (P=3.01×10^−16^). None of the other functional categories were significantly enriched for the heritability of income. When examining cell-type specific enrichment using partitioned heritability we show, for the first time, that the greatest level of enrichment for cell type specific groupings comes from the brain and central nervous system as this grouping contained 14.9% of the SNPs which collectively accounted for 43.4% of the heritability. This enrichment was also significant (P=1.02×10^−11^). Significant enrichment was also found for the cell-specific adrenal/pancreas tissues (P=0.002), cardiovascular tissues (P=0.001), and skeletal muscle tissues (P=0.002) (Figure 5A, Figure 5B, & **Supplementary Table 14**).

**Figure 5A.**
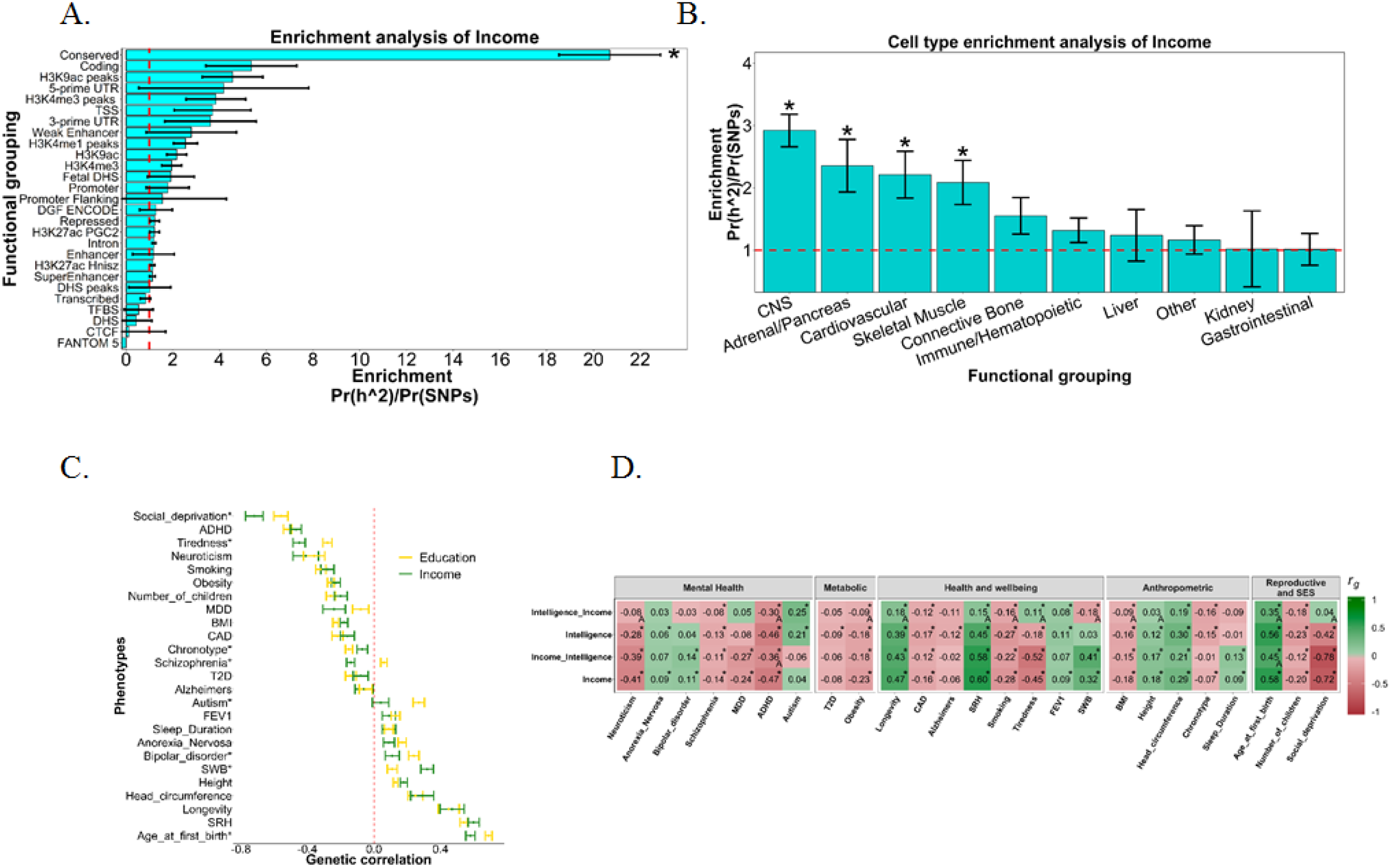
Enrichment analysis for income using the 52 functional categories (27 categories describing enrichment within these categories is shown. The full results including the functional categories with an additional 500kb boundary can be found in **Supplementary Table 14**). This analysis differs from that presented in Figure 3 and Figure 4 as here all SNPs are used, not only those that reached genome wide significance (Figure 3) or SNPs that were located within protein coding genes (Figure 4). The enrichment statistic is the proportion of heritability found in each functional group divided by the proportion of SNPS in each group (Pr(h^2^)/Pr(SNPs)). The dashed line indicates no enrichment found when Pr(h^2^)/Pr(SNPs) = 1. A Bonferroni correction controlling for 52 tests was used to ascertain statistical significance which is indicated by an asterisk. **Figure 5B**. Cell type enrichment analysis of income was performed using all SNPs as per **Figure 5A**. Here, enrichment of heritability for income was examined in 10 tissue types. A Bonferroni correction was used to control for the number of independent tests conducted, and an asterisk indicates statistical significance. **Figure 5C**. Genetic correlations between income and education with 26 phenotypes are compared. In instances where the genetic correlations were significantly different between income and education using a two-sided test (2*pnorm(-abs(abs(r_gi −_ r_gj_) / sqrt(SE_i_^2+SE_j_^2)))) an asterisk is seen by the phenotype label. Abbreviations, MDD, major depressive disorder; ADHD, attention deficit hyperactivity disorder; T2D, type 2 diabetes; CAD, coronary artery disease; SRH, self-rated health; SWB, subjective wellbeing; BMI, body mass index. **Figure 5D**. Heatmap showing the genetic correlations derived using income, intelligence as well as income conditioned on intelligence, and intelligence conditioned on income using mtCOJO. Colour indicates the direction of the genetic correlation whereas shading indicates its magnitude. Control for multiple testing was performed using FDR conducted for each of the four phenotypes separately. An asterisk indicates statistical significance of the genetic correlation. An ‘A’ indicates that there was a significant difference between the intelligence and intelligence conditioned on income (intelligence_income), or between income and income conditioned on intelligence (intelligence_income) using a two tailed test (2*pnorm(−abs(abs(r_gi −_ r_gj_) / sqrt(SE_i_^2+SE_j_^2)))).

### Causal links with intelligence

Mendelian randomization was performed using the genetic instrument derived using 19 SNPs associated with intelligence from a meta-analysis of a GWAS of intelligence from the INTERVAL BioResource^47, 48^ as well as publicly-available sources (**Supplementary data**). Here we identified a strong, causal link between intelligence and income (Beta=0.213, SE=0.063, P=7.63×10^−4^) (**Supplementary Table 15**). This indicates that greater intelligence causes a higher level of income. Sensitivity analyses revealed little evidence of directional pleiotropy which can bias the result of an MR analysis (Intercept=0.010, SE=0.007, P=0.189) (**Supplementary Table 15**). Of note is that the heterogeneity statistics (**Supplementary Table 15**) indicate that the magnitude of the causal effect is inconsistent across the SNPs used to create the instrument. However, since there was no evidence of directional pleiotropy, the overall causal estimate based on all of the genetic variants is unlikely to be biased.

### Genetic correlations

Next, we calculated genetic correlations between household income and a set of 26 data sets covering psychological traits, mental health, health and wellbeing, anthropometric traits, metabolic traits, and reproduction.

First, we build on the findings of Hill et al. (2016)^7^ by using a larger, better-powered dataset on income to show that the genetic variants associated with household income are linked with those that influence cognitive abilities, with strong genetic correlations found (intelligence, r_g_=0.69, SE=0.02, P>10×10^−200^, education, r_g_ = 0.78, SE=0.01, P>10×10^−200^). With this larger data set we show there are genetic correlations between income with health (self-rated health, r_g_=0.60, SE=0.03, P=5.72×10^−73^), mental health (subjective wellbeing, r_g_=0.32, SE=0.04, P=4.99×10^−17^), as well as longevity (r_g_=0.47, SE=0.07, P=1.29 × 10^−10^). Novel genetic correlations were identified between income and age of first birth (r_g_=0.58, SE=0.03, P=8.81×10^−99^) and the number of offspring (r_g_=−0.20, SE=0.04, P=8.81×10^−7^) as well as feelings of tiredness and fatigue (r_g_=−0.45, SE=0.04, P=3.70×10^−34^). Together, these findings indicate that genetic variants that are associated with higher income are also correlated with the genetic predisposition for a greater level of intelligence and education, a longer lifespan, better physical and mental health, fewer feelings of tiredness, having fewer children, and having better living conditions. It should, however, be noted that income shows a positive genetic correlation with the mental health variables of anorexia nervosa (r_g_=0.09, SE=0.03, P=9.53×10^−3^) and bipolar disorder (r_g_=0.11, SE=0.04, P=1.20×10^−2^) (Figure 5C & **Supplementary Table 16**).

Second, as SEP is a multi-dimensional construct and each marker of SES is imperfectly correlated with the others, the magnitude of the genetic correlations derived using income were compared with those derived using another measure of SEP, educational attainment. The goal of these analyses was to indicate if the genetic associations between household income with health differed from those of education with health. As can be seen in Figure 5C, whereas the magnitude and direction of the genetic correlations derived using income and EA with the 26 health and wellbeing, anthropometric, mental health, and metabolic traits were highly similar, there were instances of divergence indicating unique genetic associations with the two SEP variables. Of note are the variables of autism and schizophrenia. As found in previous studies^8, 49, 51, 65, 66^ schizophrenia showed a small positive genetic correlation with EA (r_g_=0.06, SE=0.02, P=1.15 × 10^−3^) whereas, in the present study, income showed a negative genetic correlation with schizophrenia (r_g_=−0.14, SE=0.02, P=6.49×10^−9^); the difference between these two genetic correlations was significant (P=6.57×10^−11^). Autism also showed a positive genetic correlation with EA (r_g_=0.27, SE = 0.03, P=1.10×10^−15^) as found previously,^8, 51, 67^ whereas income showed no detectable genetic correlation with autism (r_g_=0.04, SE=0.05, P=0.37), and this difference was again significant (P=1.17×10^−11^). Six other traits showed significantly different genetic correlations when comparing those derived using income with those derived using EA (subjective wellbeing, P=1.42×10^−5^, tiredness, P=1.60×10^−4^, age at first birth, P=1.24×10^−3^, bipolar disorder, P=1.41×10^−2^, social deprivation, P=1.72×10^−2^, and chronotype, P=3.83×10^−2^) (Figure 5C & **Supplementary Table 16)**.

Third, the role of intelligence in mediating the effect of genetic variation on income was explored by estimating the genetic correlation of income with each of the traits after conditioning the income GWAS on a GWAS on intelligence. As can be seen in Figure 5D, after controlling for intelligence the genetic correlations between income and the 26 health and wellbeing, anthropometric, mental health, and metabolic traits remained largely similar. Two exceptions to this were age at first birth, where the genetic correlation with income decreased from r_g_=0.58 (SE=0.03, P=8.81×10^−99^) to r_g_=0.45 (SE=0.04, P=1.20×10^−35^) (P_diff_=0.004), and ADHD which decreased from r_g_=−0.48 (SE=0.03, P=2.20×10^−45^) to r_g_=−0.36 (SE=0.04, P=1.86×10^−17^). This means that the genetic variation that is associated with income, but not intelligence, shows as much overlap with the 26 traits used here, as the genetic variation that is common to both income and intelligence.

In contrast, 12 genetic correlations with intelligence changed after controlling for income. Subjective wellbeing showed no genetic correlation with intelligence (r_g_=0.03, SE=0.03, P=0.31), as previously found^8^; however, subjective wellbeing was negatively genetically correlated after adjusting for income (r_g_=−0.18, SE=0.04, P=3.11×10^−5^), (P_diff_=9.92×10^−5^). The genetic correlation between intelligence and social deprivation (as measured by Townsend Scores) of r_g_=−0.42 (SE=0.04, P=1.38×10^−23^), attenuated to r_g_= 0.04 (SE=0.05, P=0.38); (P_diff_ =1.29×10^−13^). The genetic correlation between intelligence and neuroticism^8, 68^ (r_g_=−0.28, SE=0.07, P=1.75×10^−5^) also attenuated to close to zero after conditioning on income (r_g_=−0.08, SE=0.07, P=0.22), (P_diff_=0.039). This means that the genetic variation that is associated with intelligence, but not income, shows less overlap with the 26 traits used here, than the genetic variation that is common to both intelligence and income. Significant attenuations towards zero were also observed for the genetic correlations derived using intelligence once conditioning on income for the variables of self-rated health (P_diff_=6.76×10^−12^), age at first birth (P_diff_=1.33×10^−8^), fatigue or tiredness (P_diff_=6.82×10^−8^), ADHD (P_diff_=5.55×10^−4^), height (P_diff_=2.59×10^−4^), BMI (P_diff_=0.013), obesity (P_diff_=1.60 × 10^−2^), longevity (P_diff_=0.014), smoking (P_diff_=0.032) (Figure 5D, **Supplementary Table 16**).

### Genetic prediction

Polygenic risk scores were derived using the summary statistics from our GWAS of household income and the GS:SFHS data on household income. When examining the polygenic risk scores within each of the five income groups in GS:SFHS we found those in category 5 (those earning more than £70,000) had the highest PGR scores (Figure 6A). The predicted income for the PGR scores was lower in each subsequent level of household income in GS:SFHS.

**Figure 6A.**
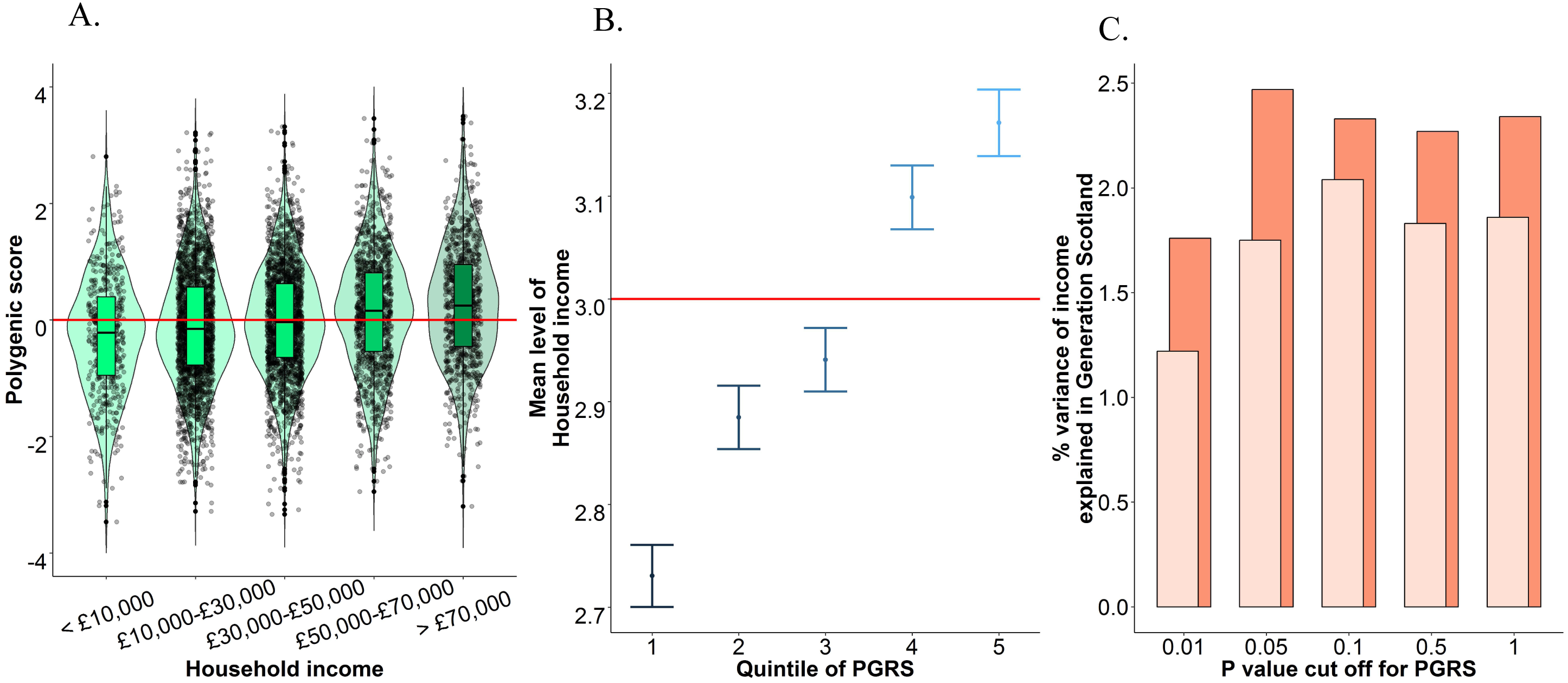
Violin plot showing the level of household income in GS:SFHS plotted against the standardized polygenic score of income in each group. Median and interquartile range are plotted. Summary data from the income GWAS performed in UK Biobank was used to derive PGRSs. Red line indicates a standardized polygenic score of 0. **Figure 6B**. The average level of household income for the five PGRSs is shown. Summary data from the income GWAS performed in UK Biobank was used to derive PGRSs. The y-axis corresponds to the 5 point classification of household income in Generation Scotland. Above the red line indicates a level of household income between £30,000 and £50,000 and below indicates a level of household income between £10,000 and £30,000 in Generation Scotland. **Figure 6C**. The variance explained by each of the five P-value cut offs for the PGRS. Light orange indicates that the income phenotype derived in UK Biobank was used to generate polygenic risk scores along with Generation Scotland. The dark orange bars indicate instances of where the MTAG phenotype derived using income and educational attainment was used to derive polygenic risk scores in Generation Scotland. Summary data from the income GWAS performed in UK Biobank, and the MTAG analysis of income was used to derive PGRSs. All results can be found in **Supplementary Table 17**.

Those in the lowest quintile of the polygenic score for income were found on average to have the lowest predicted income (Figure 6B) with the mean level of household income rising across each quintile. Those in the three lowest quintiles for their genetic propensity for income were found to have an average level of household income between £10,000 and £30,000, whereas those in the top two quintiles were found to have a household income of between £30,000 and £50,000. Polygenic prediction conducted using the summary data from UK Biobank applied to the GS:SFHS data showed that between 1.2% and 2.0% of the variance in household income can be predicted using the polygenic score for income (**Supplementary Table 17** & Figure 6C) with the PGRS that was most predictive using a P-value cut off of 0.1.

### Multi-trait analysis of genome-wide association studies

MTAG has previously been used to conduct the first well-powered GWAS on intelligence.^8^ We used MTAG here to increase the power of our GWAS on income by meta-analysing it with another measure of SEP, educational attainment^10^ as measured by the number of years of education a participant has completed. MTAG was conducted using the default settings and applied to increase the power in the GWAS of household income. Following the application of MTAG, the mean χ^2^ statistic increased from 1.45 to 1.73 and increased the effective sample size to 505,541 for income. The maxFDR was calculated to determine the credibility of these SNPs as being trait-specific to income. The maxFDR derived was 0.003, over an order of magnitude lower than the commonly accepted standard of false discovery and comparable with those reported previously.^8, 20^ Of note is that this FDR is trait-specific, meaning that if these SNPs were associated with EA, but not with income, then an inflation of the maxFDR would be seen. In addition, we examined the genetic correlations between the MTAG-income phenotype with the income from a previous GWAS.^7^ We find that the genetic correlation between our MTAG-income phenotype and a previous GWAS on income^7^ was r_g_=0.97 (SE=0.024), with a genetic correlation of r_g_=0.94 (SE=0.004) with educational attainment. This indicates that the polygenic signal in the MTAG-income analysis is virtually identical to that found in previous GWAS of income, but also that it captures more of the variance that is shared between income and education.

The heritability of the MTAG-income phenotype was 8.1% (SE = 0.3%) and the LDSC regression intercept was 0.92 (SE = 0.98) indicating that the increase in mean χ^2^ was not due to capturing the effects of stratification, cryptic relatedness or other confounds, but rather due to a more accurate estimate of the polygenic signal in the data.

Using this MTAG-income phenotype we identify 144 independent genomic risk loci (**Supplementary Figure 2A** & **Supplementary Table 18**). A total of 24 of these loci overlapped with the 30 identified without using MTAG meaning that by using MTAG an additional 120 independent loci were identified that were associated with income (**Supplementary Table 19**). Functional annotation of these loci, as well as gene-based analyses and partitioned heritability analysis showed results that were consistent with a better-powered GWAS dataset on household income. These results can be found in the supplementary results section.

Polygenic risk scores analysis was also performed using the MTAG derived income phenotype where the accuracy of phenotype prediction was increased compared to using the GWAS on income alone, as would be theoretically expected.^20^ Using the MTAG phenotype, between 1.7% and 2.5% the variance of income was predicted in an independent sample (**Supplementary Table 17** & Figure 6C) with the PGRS that was most predictive using a P-value cut off of 0.05. The lower threshold being more predictive is indicative of the greater power of the MTAG data set relative to the data set derived using income alone, as the power reliably to detect small effects is greater.

## Discussion

Using the UK Biobank data set, we identified 30 independent genetic loci associated with income levels in Great Britain today. These loci collectively harbored 3,712 SNPs that attained genome-wide significance, representing a considerable advance on the two loci previously identified by Hill et al. (2016).^7^ Using these data, we were able to make a number of novel findings that highlight the genetic contributions to a marker of socio-economic position.

First, the loci identified with income showed clear evidence of functionality, particularly regarding their links to gene expression, regulatory regions of the genome, and open chromatin states. Second, by combining our GWAS data with eQTL data from BRAINEAC,^69^ GTEx,^70^ and others, along with chromatin interaction data^71, 72^ we were able to prioritize which genes were likely to be causal based on the overlap of multiple lines of biological enquiry. Although income, as a biologically distal phenotype, will not be linked to genetic variation directly (Figure 1)^7^, genes that may exert a causal influence are likely to do so through their effect on more proximal phenotypes.^12^ It is important to note that not all of the implicated genes are certain to play a causal role in income differences. The analyses conducted here serve to identify which genes are potentially informative as to the underlying biology of these differences.

Using our GWAS data set on household income, we identified 47 genes that were mapped to the 30 independent genomic loci using positional, eQTL, and chromatin mapping. In addition, we used the 118 genome-wide significant genes from our gene-based analysis of income to further refine this set to a total of 24 implicated genes. These 24 genes therefore should be prioritized in follow-up studies as they are located close to the associated loci, have expression correlated with genetic variation of the SNPs in the independent genomic loci, have chromatin interactions taking place between these genes and the SNPs found in the independent loci (**Supplementary Table 5**) and, consistent with highly polygenic traits, these genes harbor many SNPs that show consistent associations with income (**Supplementary Table 7**). In addition, 18 of these genes have been associated with intelligence^8^, so efforts to ascertain how such genetic variation is associated with income differences should examine their associations with intelligence more closely.

Third, by broadening our analysis to include the polygenic signal present in our data that fell outside of the independent loci, we identified additional, novel, functional elements of the genome linked to differences in income. By combining the gene-based statistics from MAGMA with gene expression data from the GTEx^70^ database, we identified a positive association between expression in the brain, as well as several specific regions, and the level of association displayed by the gene-based statistics on income. This indicates that the higher the level of association between a gene and income, the higher that gene’s level of expression in the brain will be.

Cell type specific analysis revealed that the expression that was specific to the serotonergic neurons and to medium spiny neurons was associated with income. Medium spiny neurons have previously been linked to schizophrenia^42^ as well as to education.^10^ Both schizophrenia and education have strong cognitive components and have previously been linked to glutamatergic systems including the NMDA receptor signaling complex.^73^ Medium spiny neurons are, however, a sub-type of GABAergic inhibitory neurons. Future work should examine if, like other cognitive traits, income is linked to both GABAergic and glutamatergic systems.

Using partitioned heritability analysis, we found that conserved regions of the genome were enriched for genetic associations with income. These regions contained 2.6% of the SNPs but these collectively accounted for 53.9% of the SNP heritability of income. These regions are subjected to ongoing purifying selection and, under models of neutral selective pressure, accumulate base-pair substations at a lower rate than SNPs drawn from outside of these regions. This finding may indicate the presence of a mutation-selection balance acting on the partly-heritable phenotypes that contribute towards income differences. A mutation-selection balance is characterized by the removal of deleterious variants from a population at the same rate that novel mutations occur.^74^ Also consistent with the action of a mutation-selection balance was the observed correlation of 0.42 (P=2.2 × 10^−4^) between minor allele frequency and the effect size for the SNPs found in the independent genomic loci – this indicated that variants with a lower MAF have a greater association with income.

Partitioned heritability analysis also identified enrichment across multiple cell types, indicating that a wide range of tissues types may be associated with phenotypes that are associated with income differences. Whereas the tissue of the central nervous system showed the highest level of enrichment, the adrenal/pancreas, skeletal muscle, cardiovascular, and immune/hematopoietic tissues all showed significant enrichment. The finding that the regions of the genome undergoing purifying selection, as well as tissue types from multiple biological systems are enriched in their associations with income, is consistent with the notion that, whereas intelligence differences might make some contributions to differences in income, a range of other partly-heritable phenotypes also likely to contribute. This might include susceptibility to some diseases, as evidenced by the disparate tissue types linked to income.

These two approaches, gene-based statistics and LDSC regression, illustrate how combining the genetic data from GWAS with gene expression data can be informative as to the possible biological processes that are associated with income. This is of particular value for traits, like income, that are biologically distal and, as such, have no clear biological analogue. This combination of data provides evidence that some of the individual differences in income are related to the genetic basis of gene expression differences in the brain (Figure 4B), as well as highlighting the role of specific classes of neuron (Figure 4C). As importantly, we show the role for tissue types outside of the central nervous system (Figure 5B) indicating that genetic factors associated with income differences also lie outside of the phenotype of intelligence, and outside of cortical tissue types.

Fourth, using Mendelian Randomization, we provided the first evidence implicating intelligence as one of the causal, partly-heritable, phenotypes that might be one bridge in the gap between molecular genetic inheritance and phenotypic consequence. This result helps explain why individual differences in income are found to be partly heritable. It is because the SNPs associated with income are likely to be mediated by other more biologically-proximal phenotypes (e.g. neuronal function etc.). This finding indicates that genetic variants that are predictive of income might in part be so because they influence intelligence, and that it is intelligence that is an arguably more proximal causal factor, accounting for some of the differences in income in Great Britain today.

Fifth, we provide the best-to-date estimates of genetic correlations with income showing a genetic correlation with longevity (r_g_=0.47, SE=0.07, P=1.29×10^−10^). More importantly, this paper used genetic correlations to determine if the genetic contributions to income were significantly different in relation to health than another measure of SEP, years of education. The estimate of these significantly different genetic correlations indicates that, relative to the genetic underpinnings of education, the genetic variants that associated with higher income are also associated with lower levels of social deprivation, fatigue or tiredness, schizophrenia, with no increased risk of autism, in addition to a greater level of subjective wellbeing, and a lower age at which an individual has their first child (Figure 5C).

In summary, these results indicate that different markers of SEP have genetic contributions that are differentially associated with health. Specifically, whereas the genetic underpinnings of education and income are very similar, in instances where they do differ, it is the genetic associations with income that are more protective and less harmful for physical and mental illness than the genetic associations with education.

These significantly different genetic correlations with education and income may also indicate that educational attainment serves to provide access to opportunities in the labor market, and those that have these opportunities are then better placed to engage in health-relevant behaviors. This would indicate that, whereas income may be a more distal phenotype from DNA than education, it is closer to outcomes such as later-life health, as shown by the significantly different genetic correlations. Future work should examine models where DNA > neuronal properties -> intelligence -> education -> income -> health, using multivariable Mendelian randomization^75–77^ to gauge the direct and indirect effects of income and education on health outcomes.

We found that, when the genetic associations that are shared between income and intelligence were removed, leaving only the genetic associations that are specific to income, the genetic correlations with other traits are largely unchanged. The exceptions were with ADHD and with age of first birth, where the genetic correlations with income are both significantly attenuated once conditioned on intelligence. However, by conditioning intelligence on income, there was a significant change in the magnitude of 12 of the genetic correlations. These results indicate that the genetic variation associated with intelligence and income is also associated with many health and mental health traits, because, when this shared variance is removed, leaving only the variance that is unique to intelligence, the magnitude of the genetic link between intelligence and health is reduced. In the case of the genetic link between intelligence, social deprivation, neuroticism, and height, this genetic association disappears entirely. The exception is that subjective wellbeing as a genetic correlate of intelligence is found only after the variance that is common to both income and intelligence is removed.

One interpretation of this finding is that the residual variance left in income after conditioning on intelligence still contains the genetic contributions to other partly-heritable traits (such as conscientiousness, or resistance to disease). These traits also contribute towards individual differences in income and so the association between income and health is, largely, intact following conditioning on intelligence. This would imply that intelligence is only one of a number of factors that contributes to variation in income, but income is a very important factor that mediates the associations between intelligence and health. Future work examining the genetic relationship between income and health, as well as intelligence and health, should focus on this genetic overlap between intelligence and income using tools such as genomic structural equation modelling (SEM) to partition the total variance of traits like income into the variance that is shared with intelligence and the variance that is separate from it.^78^ Sixth, we used our income-based GWAS derived in UK Biobank, along with data from Generation Scotland, to create polygenic scores that were used to predict income. We were able to predict between up to 2.00% of variance in income using DNA alone into an independent sample. Previous estimates of income prediction using DNA have been able predict 0.8% of the variance. Our results show that even for phenotypes that are not impacted directly by genetic effects, but rather are more biologically distal as is the case with income, that the link between genotype and phenotype is sufficient to make meaningful predictions, based on DNA alone.

Seventh, using MTAG, we were able meta-analyze our GWAS on household income with a GWAS on educational attainment to add power to our GWAS on income. The associations generated in this way were specific to income, and we increased our effective sample size from 286,301 participants to 505,541. This increase in power was accompanied by an increase in the loci associated with income that increased from 30, using income alone, to 144 once meta-analyzed using MTAG. Of these 144 associations, 120 of were not found to be genome-wide significant before the application of MTAG. These loci demonstrated the same patterns of functional enrichment as shown in the 30 loci identified using income alone. We also identified the same relationship between expression in the brain, and across multiple cortical structures, using the better powered MTAG-derived income phenotype (Supplementary results). Furthermore, following meta-analysis with MTAG, we were able to increase our prediction accuracy of income by 25%.

The limitations of this study include that income was measured at the level of the household, and was not an individual-level measure of income. However, previous GWASs examining household income variables have shown that income, measured on a household level, has a genetic correlation of 0.90 (S.E. = 0.04) with educational attainment, as measured on an individual level, indicating that the household-level effects might be generalizable to individual persons. Furthermore, GWASs conducted on regional measures of educational attainment show genetic correlations of >0.9 with education measured using an individual’s own level of educational attainment.^79^ A limitation of the Mendelian Randomization analysis specifically is that the estimates may be due to dynastic effects, whereby genes from the parent are associated with parental behaviors, which are a causal factor in the SES of the child.^80^ An example of this would could be that parents with a greater predisposition towards intelligence are also those that are more likely to provide opportunities for their children to enter higher-income occupations. Whereas the current data cannot differentiate between causality and dynastic effects, it should be noted that, for another measure of SEP, educational attainment, whereas there are indirect genetic effects, these account for almost half of the variance of the direct genetic associations.^61^ Future work in multi-generational samples should examine the role that such indirect genetic effects play in individual differences in income, as well as if their presence (if established) could result in an inflation of the estimate for a causal effect using Mendelian Randomization.

Another limitation is that the present study was restricted to examining common genetic effects. Should rare or less common genetic variation be associated with income, then these effects will be absent from this study. Future work should utilize methods that can capture these genetic effects,^81^ as well as examine SEP variables using whole exome or whole genome sequencing. In addition, the participants of UK Biobank are drawn from the more educated individuals in the UK, which might introduce collider bias.^82^ Whereas a comparison of the level of SEP between the individuals in UK Biobank and the census conducted in the UK indicates that SEP, as measured using the Townsend Deprivation Index,^83^ was very similar,^7^ future work aiming to quantify or control for collider bias would be of value in addressing this potential issue.

Finally, it should be noted that GWASs, like heritability estimates, describe differences that exist within populations. This means that, although we report here that those with a greater number of intelligence-associated genetic variants tend to be those who report higher incomes, it does not hold that this is true across all societies or times. Indeed, the links between markers of SEP and health are not consistent across all societies.^84^ Research into genetic links to education has found indications that the genetic variants linked to higher educational attainment are less predictive of success in societies that have less meritocratic selection for education and occupation.^85^ Future work examining the relative contribution of genetic and environmental associations with income, as well as the biological systems causally implicated in any GWAS conducted on a marker of SEP across many cultures, would be valuable in identifying more and less meritocratic societies.

In conclusion, this work adds to the growing body of evidence^7^ indicating that markers of socioeconomic position, and their links to health, are not purely environmental in origin.^6^ We found that SEP variation in the Great Britain is partially accounted for by genetic differences in the population.^79^ We found little evidence that these genetic differences were attributable to population stratification, but rather that they indicated the unequal distribution of heritable traits, including intelligence, across different SEP groups. Using multiple forms of biological data, we showed that these genetic differences are predominantly found in regions of the genome that have undergone negative selection, and those linked to differences in gene expression in the brain, particularly in medium spiny neurons. We also prioritise 24 genes for further follow up as evidence from eQTL analysis, chromatin interactions, with previous associations with intelligence converging to implicate 18 of these genes. Furthermore, we identify intelligence as one of the causal psychological mechanisms partly driving differences in SEP in Great Britain today.

## Supporting information

Supplementary Results

Supplementary Figure 1A Chromosome 1

Supplementary Figure 1B Chromosome 2

Supplementary Figure 1C Chromosome 3

Supplementary Figure 1D Chromosome 4

Supplementary Figure 1E Chromosome 5

Supplementary Figure 1F Chromosome 6

Supplementary Figure 1G Chromosome 7

Supplementary Figure 1H Chromosome 9

Supplementary Figure 1I Chromosome 13

Supplementary Figure 1J Chromosome 17

Supplementary Figure 1K Chromosome 18

Supplementary Figure 1L Chromosome 19

Supplementary Figure 1M Chromosome 20

FAQ

## Acknowledgements

This work was undertaken in The University of Edinburgh Centre for Cognitive Ageing and Cognitive Epidemiology (CCACE), supported by the cross-council Lifelong Health and Wellbeing initiative (MR/K026992/1). Ethical approval for UK Biobank was received from the Research Ethics Committee (REC reference 11/NW/0382). This work was conducted under UK Biobank application 10279. Generation Scotland received core support from the Chief Scientist Office of the Scottish Government Health Directorates [CZD/16/6] and the Scottish Funding Council [HR03006]. Genotyping of the GS:SFHS samples was carried out by the Genetics Core Laboratory at the Wellcome Trust Clinical Research Facility, Edinburgh, Scotland and was funded by the Medical Research Council UK and the Wellcome Trust (Wellcome Trust Strategic Award “STratifying Resilience and Depression Longitudinally” (STRADL) Reference 104036/Z/14/Z). Funding from the Biotechnology and Biological Sciences Research Council (BBSRC), the Medical Research Council (MRC), and the University of Edinburgh and gratefully acknowledged. CCACE funding supports IJD.

Participants in the INTERVAL randomised controlled trial were recruited with the active collaboration of NHS Blood and Transplant England (www.nhsbt.nhs.uk), which has supported field work and other elements of the trial. DNA extraction and genotyping was co-funded by the National Institute for Health Research (NIHR), the NIHR BioResource (http://bioresource.nihr.ac.uk/) and the NIHR [Cambridge Biomedical Research Centre at the Cambridge University Hospitals NHS Foundation Trust] [*]. The academic coordinating centre for INTERVAL was supported by core funding from: NIHR Blood and Transplant Research Unit in Donor Health and Genomics (NIHR BTRU-2014-10024), UK Medical Research Council (MR/L003120/1), British Heart Foundation (RG/13/13/30194) and the NIHR [Cambridge Biomedical Research Centre at the Cambridge University Hospitals NHS Foundation Trust]. A complete list of the investigators and contributors to the INTERVAL trial is provided in reference^47^. The academic coordinating centre would like to thank blood donor centre staff and blood donors for participating in the INTERVAL trial.

*The views expressed are those of the authors and not necessarily those of the NHS, the NIHR or the Department of Health and Social Care.

WDH is supported by a grant from Age UK (Disconnected Mind Project). We thank George Davey Smith for his comments on an early version of this manuscript.

