## Supplementary data for "Genetic analysis identifies molecular systems and biological pathways associated with household income"

Supplemental data.

Deriving independent groups for Mendelian randomisation.

Publically available GWAS data on 78,308 individuals on whom intelligence was measured was meta-analysed with the data from the INTERVAL^1^ study. The INTERVAL^1^ study is a prospective cohort study of ~50,000 participants drawn from blood donors.^2^ Recruitment occurred between 2012 and 2014 whereby blood donors aged 18 years and older consented to be recruited from the National Health Service Blood and Transplant (NHSBT) static donor centers across England. Participants are largely healthy as individuals as eligibility for donation is contingent upon being free from major disease (myocardial infarction, stroke, cancer etc) and those who reported being unwell or having had recent illness or infection were not eligible for blood donation and so were not part of the INTERVAL sample.

Participants agreed to take part in online questionnaires that contained lifestyle and health information, including self-reported height, weight, ethnicity, current smoking status, alcohol consumption, doctor-diagnosed anemia, use of medications (hormone replacement therapy, iron supplements) and menopausal status. The INTERVAL study was approved by the Cambridge (East) Research Ethics Committee and UK Biobank was approved by the North West Multi-center Research Ethics Committee (MREC). Informed consent was obtained from all participants.

Genotyping

DNA was extracted from whole blood using the buffy coat at LGC Genomics (UK) using a Kleargene method and samples of sufficient concentration and purity were aliquoted for shipment to Affymetrix for genotyping. Standard quality control (QC) procedures were implemented by Affymetrix during the genotyping pipeline. This included the excluding samples with a poor signal intensity (dish QC <0.82) along with samples that had a low call rate (< 0.97) based on 20,000 high quality probesets. Variants were excluded in the event that they had a call rate of <0.95, has more than three clusters, which indicates of-target measurement, had cluster statistics that were indicative of poor quality genotyping or multi-allelic variants that could not be called easily (Fisher’s linear discriminant, heterozygous cluster strength, homozygote ration offset were used). A total of 48,813 participants from the INTERVAL sample were genotyped in 10 batches. Within-batch sample and variant QC was performed following standard Affymetrix QC exclusions. Non-best probesets were excluded to leave a single probeset per variant. Visual inspection of the cluster plots showed that some variants and minor allele homozygotes incorrectly called due to the presence of an extreme intensity outlier. Variants were failed from a batch in the event that there were fewer than 10 called minor allele homozygotes, the cluster plot contained at least one sample with an intensity at least twice as far from the origin as the second most extreme outlier, or if the outlying sample had an extreme polar angle (15^o^ of > 75^o^) in the direction of the minor allele.

Samples that were not of European ancestry, as well as duplicate samples, were excluded prior to further QC of the variants within each batch using a set of high quality autosomal variants. These were defined as those with a minor allele frequency (MAF) >0.05, a Hardy-Weinberg equilibrium (HWE) p value of >1×10^−6^, or if there was *r*^2^ of ≤0.2 between pairs of variants. Duplicate samples were identified as those where π̂ ≥ 0.9 using Method-of-Moments IBD approach implemented in PLINK^3^. Non-Europeans were identified as those participants who had scores on PC1 or PC2 of < 0 as derived using a principal components analysis conducted on the INTERVAL samples with the 1000 Genomes major ancestry populations.^4^ Variants that strongly deviated from HWE (p value <5×10^−6^), following a Fischer’s exact test for low-frequency and rare variants (Those with a MAF <0.05 across all ten batches) or those with a χ^2^ test for common variants were excluded from the batch. Finally, variants were also excluded from the batch in the event that they had a call rate of <0.97 (within-batch) and variants were dropped from all batches in the event that they failed in four of the ten batches due to either HWE deviation, low call rate, or failure in the Affymetrix exclusion criteria. These passing samples were then merged using the methods described by Jun et al. (2012)^5^ full details can be found in Astle et al. (2016).^6^ Non-autosomal and multi-allelic variants, and monomorphic variants were removed and the data were phased using SHAPEIT3 (<https://jmarchini.org/shapeit3/>) , to phase the data in chunks of 5,000 variants with a 250 variant overlap between chunks. Imputation was conducted using a combined 1000 Genomes Phase 3-UK10K imputation panel with imputation being carried out on the Sanger Imputation Server for the 43,059 participants that remained.

Intelligence phenotype INTERVAL

A general factor of cognitive ability was derived in INTERVAL using four tests. These were the Stroop Test (a measure of attention and reaction times), Trail Making Test B (a measure of executive function), Pairs Test (a measure of episodic memory), and a Reasoning test (a measure of intelligence). These tests have been adapted from the Cardiff Cognitive Battery^7^ and have been designed specifically for use in cognitive testing in epidemiology settings. Participants scores from each of these four tests was analysed using a principal components analysis where the first unrotated component was extracted. This component explained 48.0% of the variance in the test battery where each test loaded onto this single factor at 0.34-0.60. The effect of age (using fractional polynomials), sex and the effect of experiencing a stroke/TIA during the trial were controlled for using regression with the first unrotated principal component being the outcome variable. The residuals from this model were then used as the measure of intelligence for the 17,213 participants who had both genotype and phenotype data available.

Genotyping

Individuals from INTERVAL who also took part in UK Biobank were identified using KING^8^; heterozygote concordance rate> 80% before being removed from the INTERVAL sample. A linear-mixed model was fit using BOLT-LMM^9^ and adjusted for the first five principal components of ancestry. The final sample size was 17,213 individuals. Following association analysis variants where the minor allele frequency (MAF) was <0.001 and an r^2^ of <0.6 were removed.

Meta-analysis of INTERVAL data

Publically available summary GWAS data on 78,308^10^ participants was downloaded. These data were meta-analysed using 17,213 participants of the interval consortium using a sample size weighted meta-analysis implemented in METAL.^11^ Following meta-analysis SNPs were removed if they were not present in both the INTERVAL sample and in the publically available data set on intelligence. SNPs were also excluded in the even that MAF <0.001. Z-scores were converted into beta weights using the equation, Beta <- Z-score/sqrt(2*MAF * (1-MAF) * (N+Z-score^2)), and standard errors were derived using SE <- 1/sqrt(2*MAF*(1-MAF) * (N+Z-score^2)).^12^ This yielded a total of 10,895,565 SNPs following meta-analysis in a sample of 95,521 individuals.
