## Supplementary figures and images for "Genetic analysis identifies molecular systems and biological pathways associated with household income"

### Supplemental Figure 2

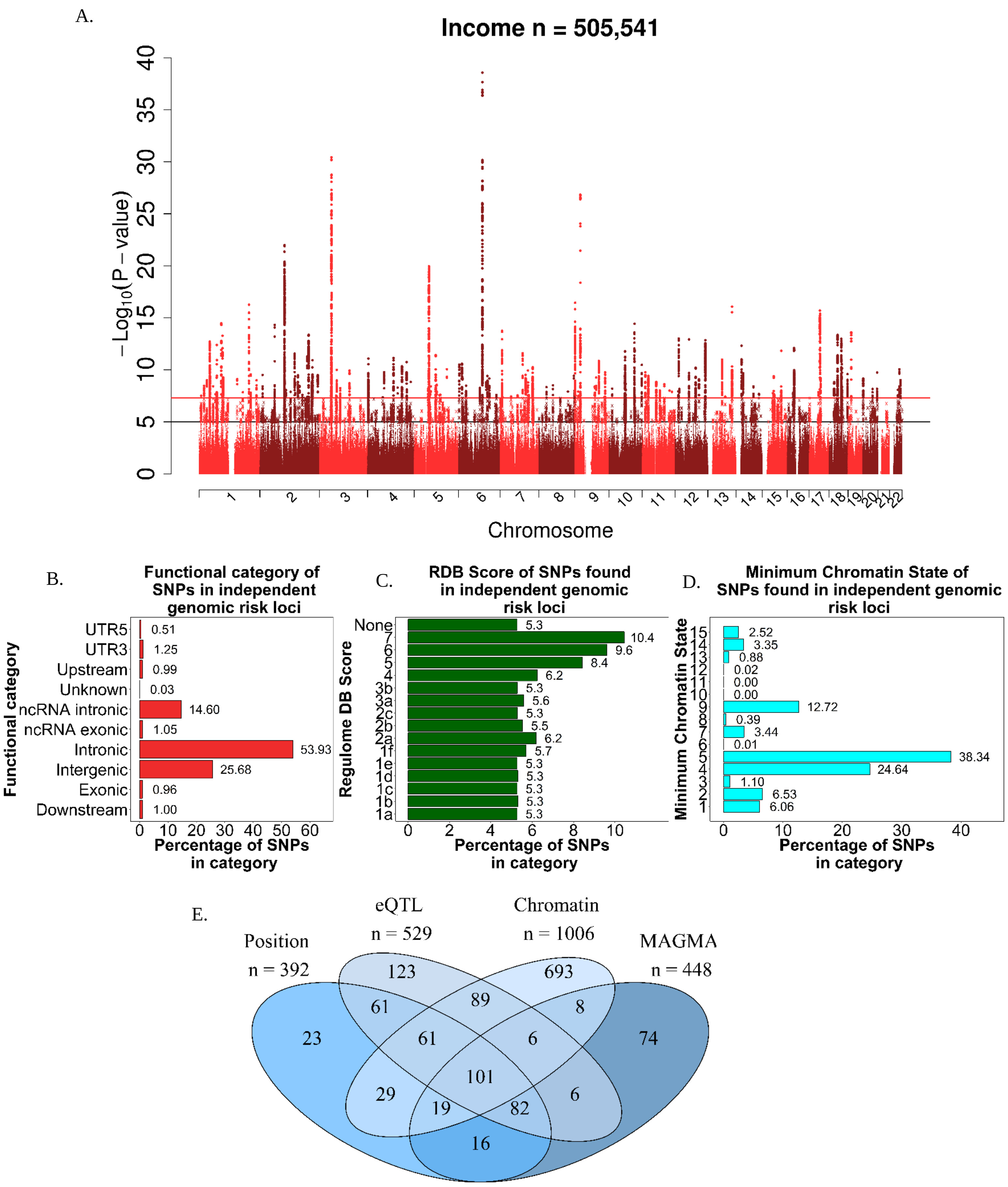

### Supplemental Figure 3

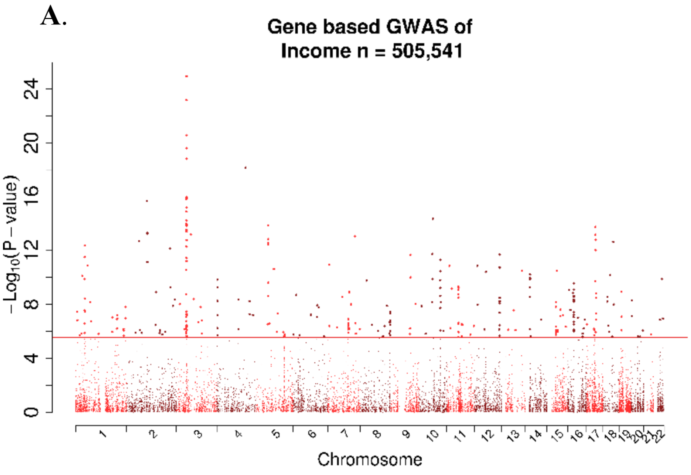

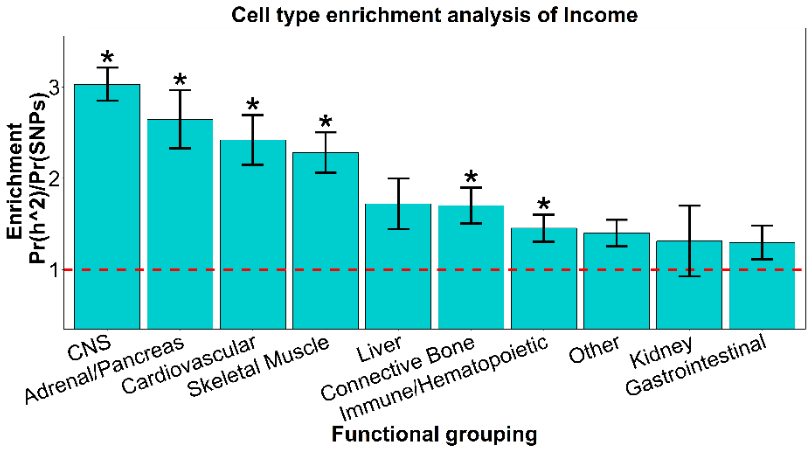

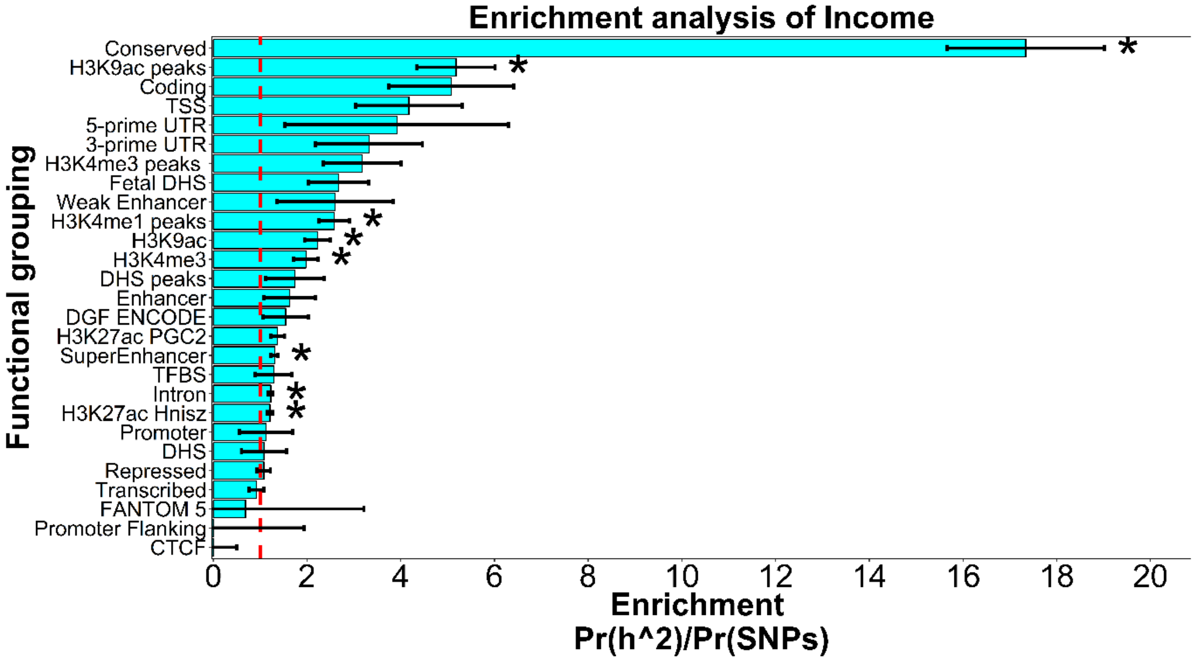

### Supplementary Figure 1A Chromosome 1

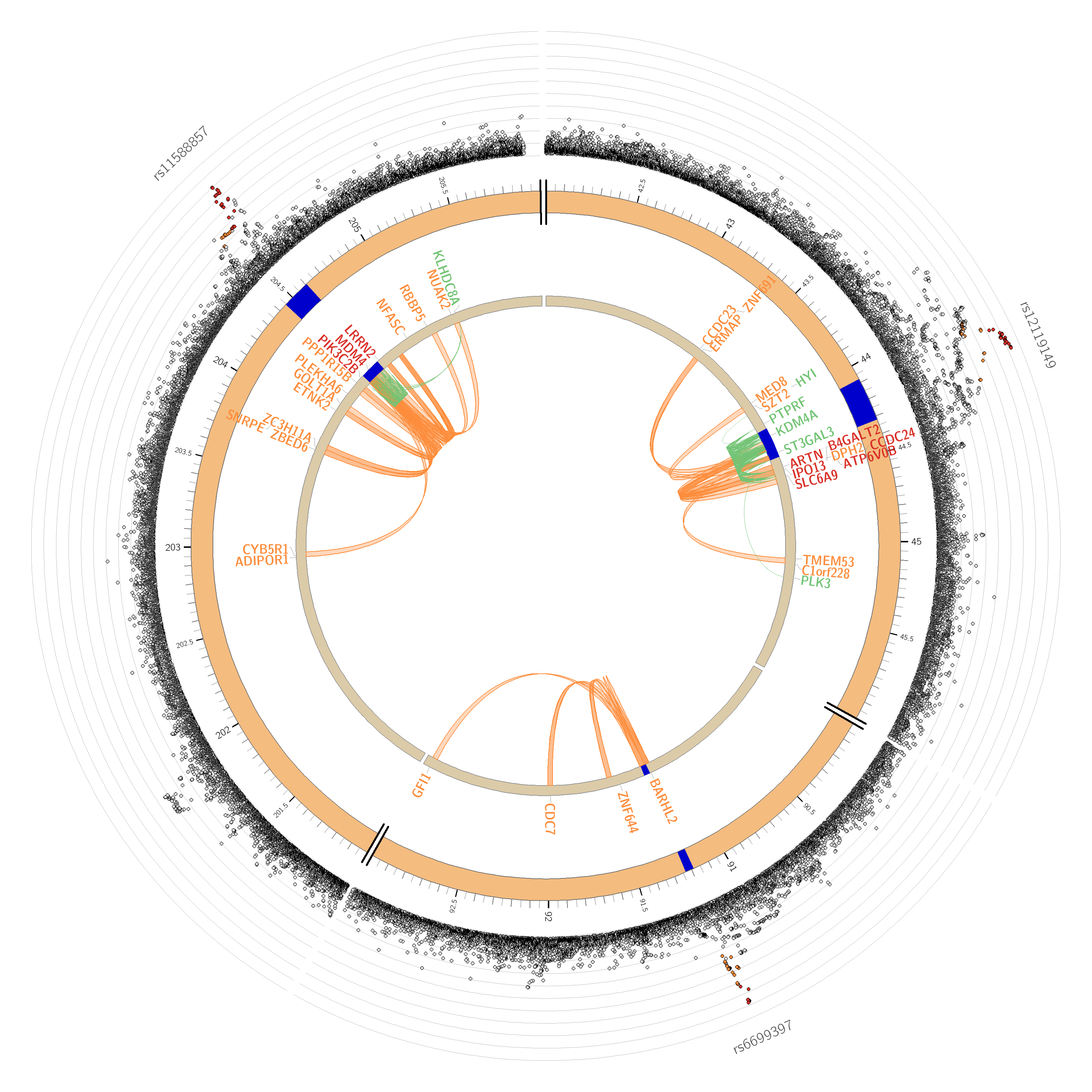

### Supplementary Figure 1B Chromosome 2

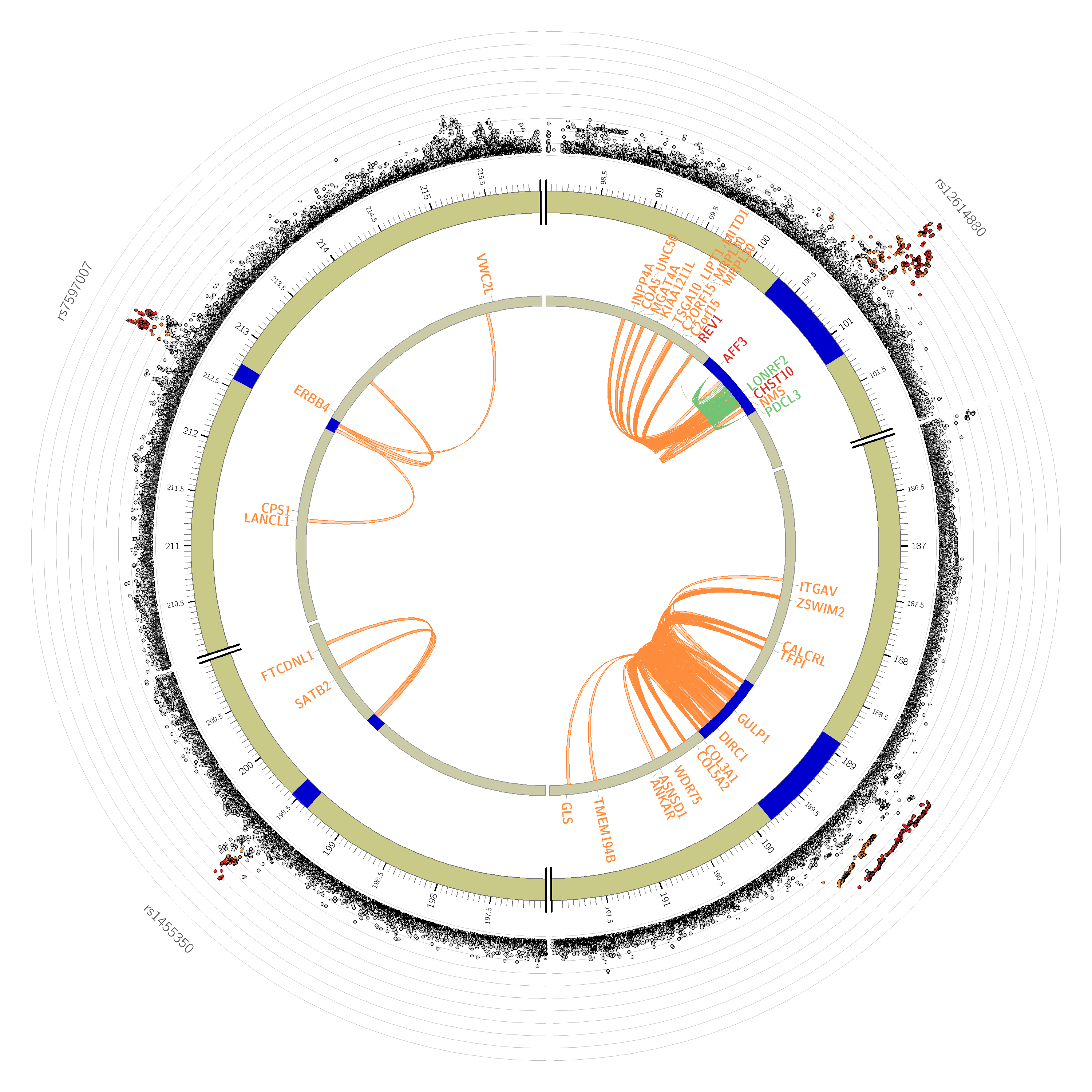

### Supplementary Figure 1C Chromosome 3

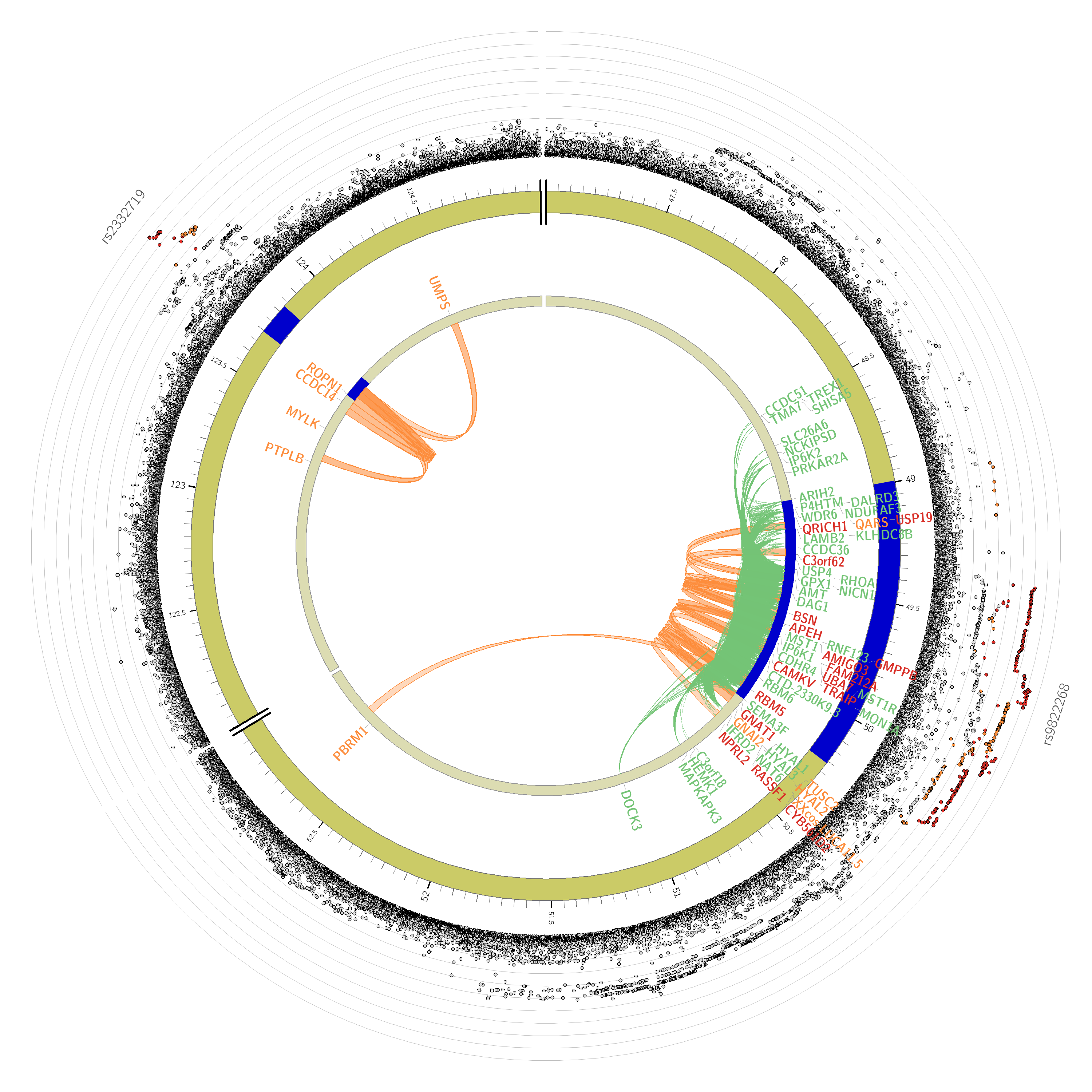

### Supplementary Figure 1D Chromosome 4

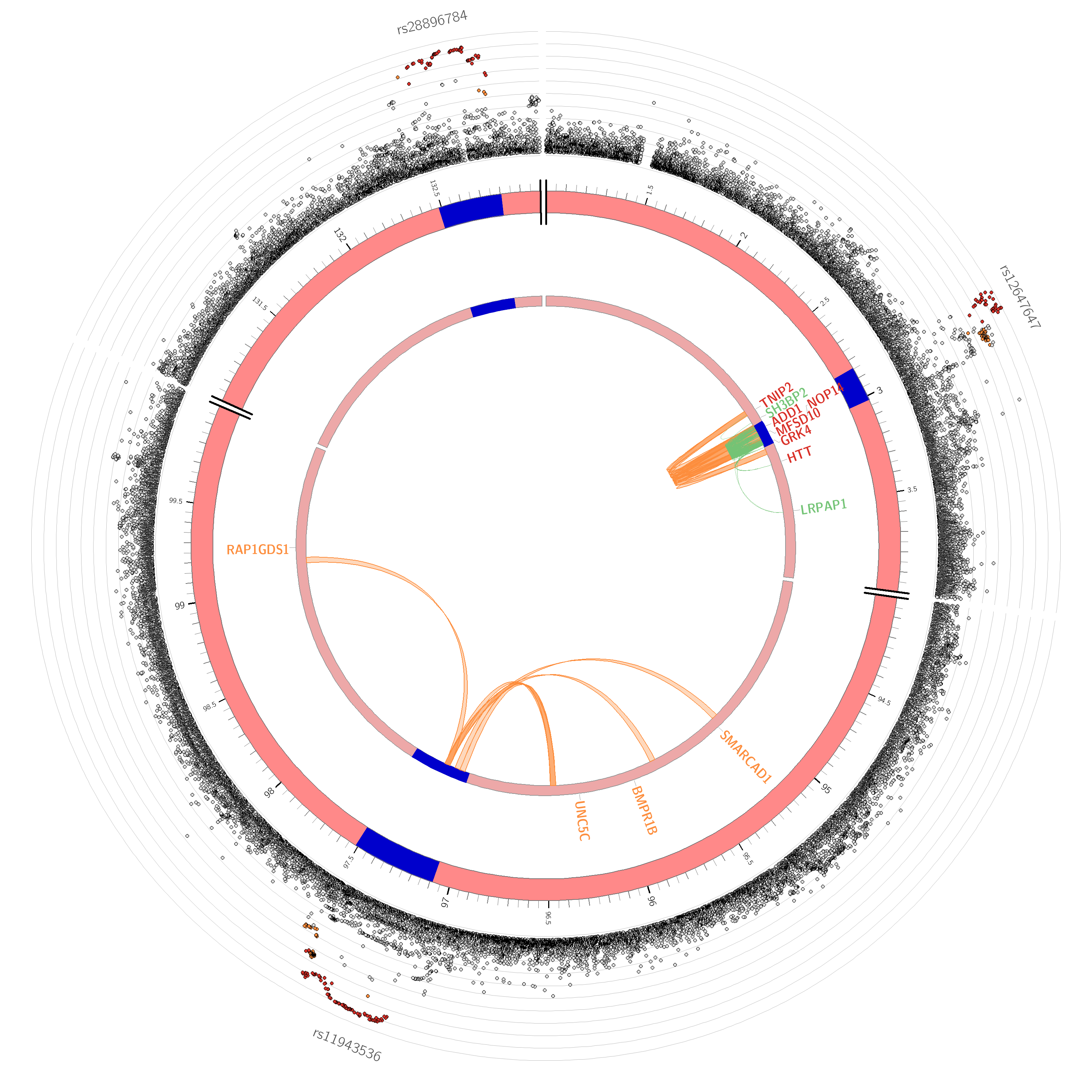

### Supplementary Figure 1E Chromosome 5

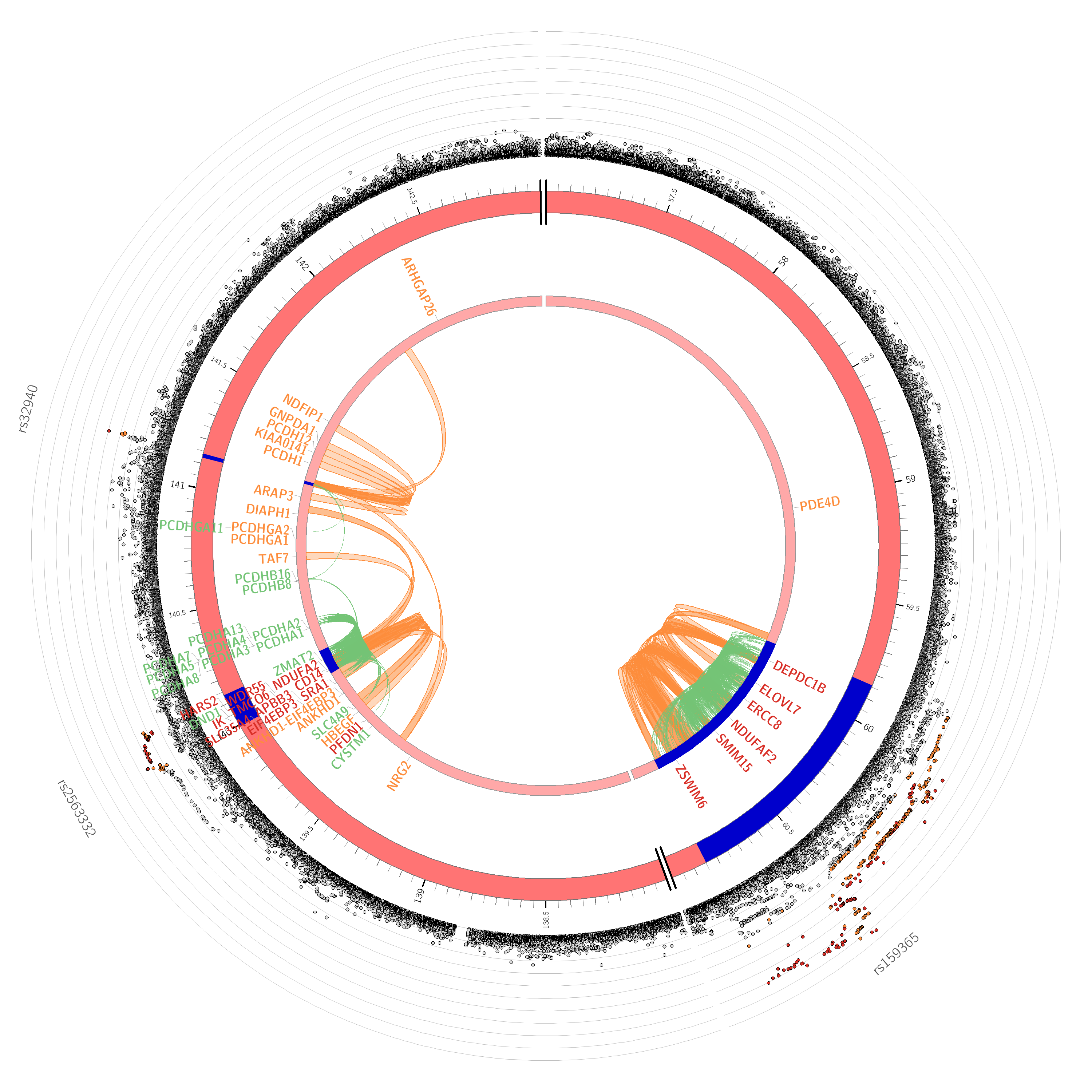

### Supplementary Figure 1F Chromosome 6

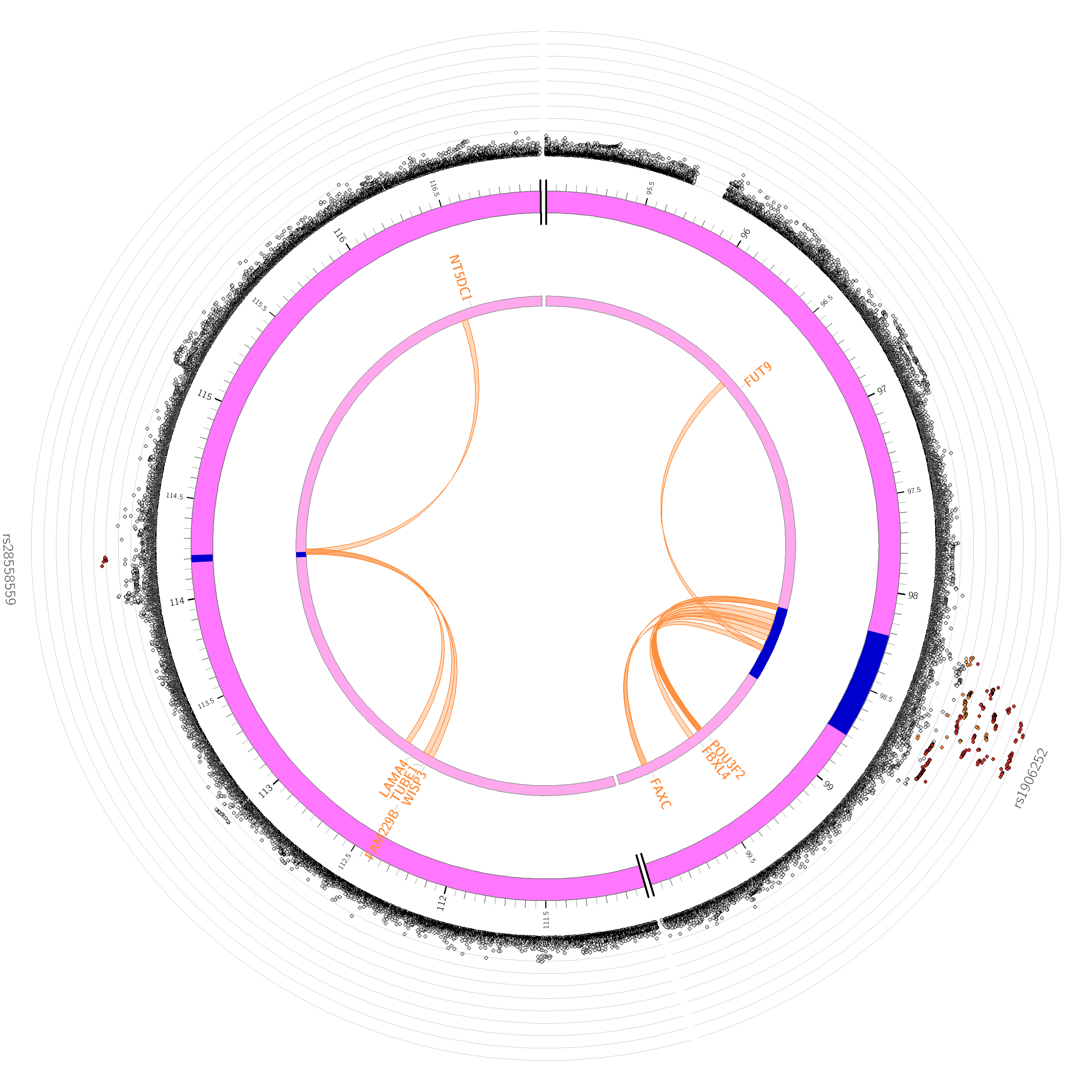

### Supplementary Figure 1G Chromosome 7

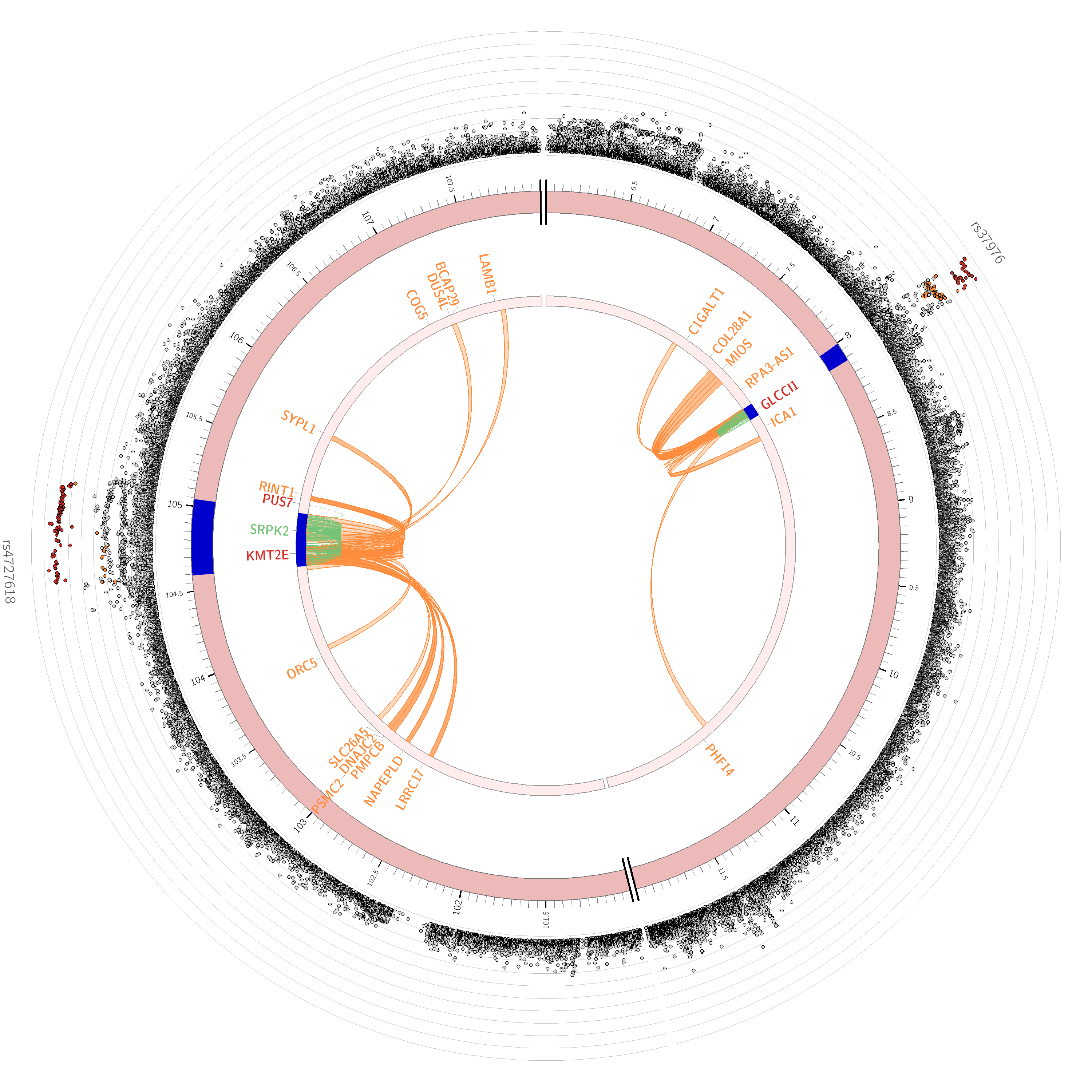

### Supplementary Figure 1H Chromosome 9

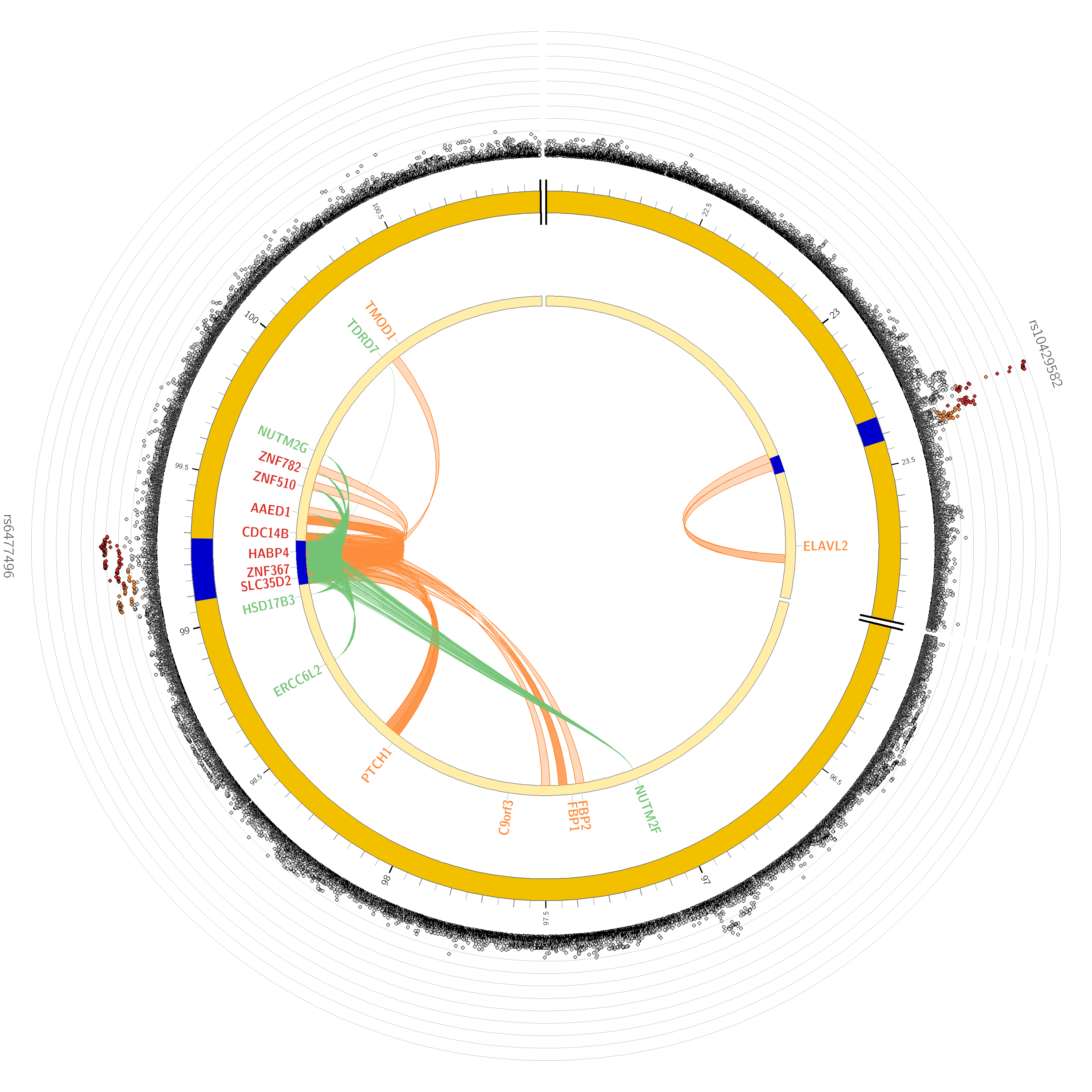

### Supplementary Figure 1I Chromosome 13

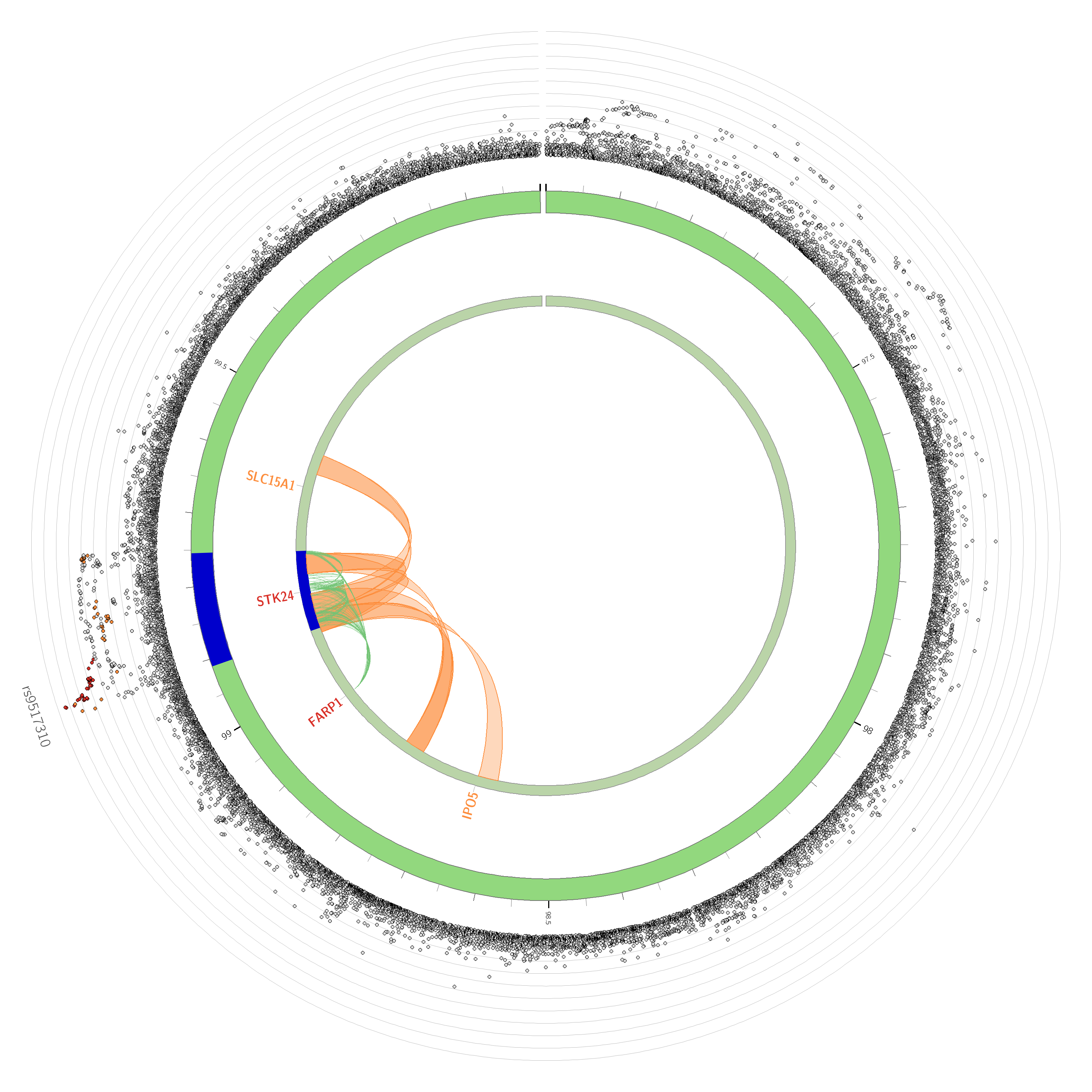

### Supplementary Figure 1J Chromosome 17

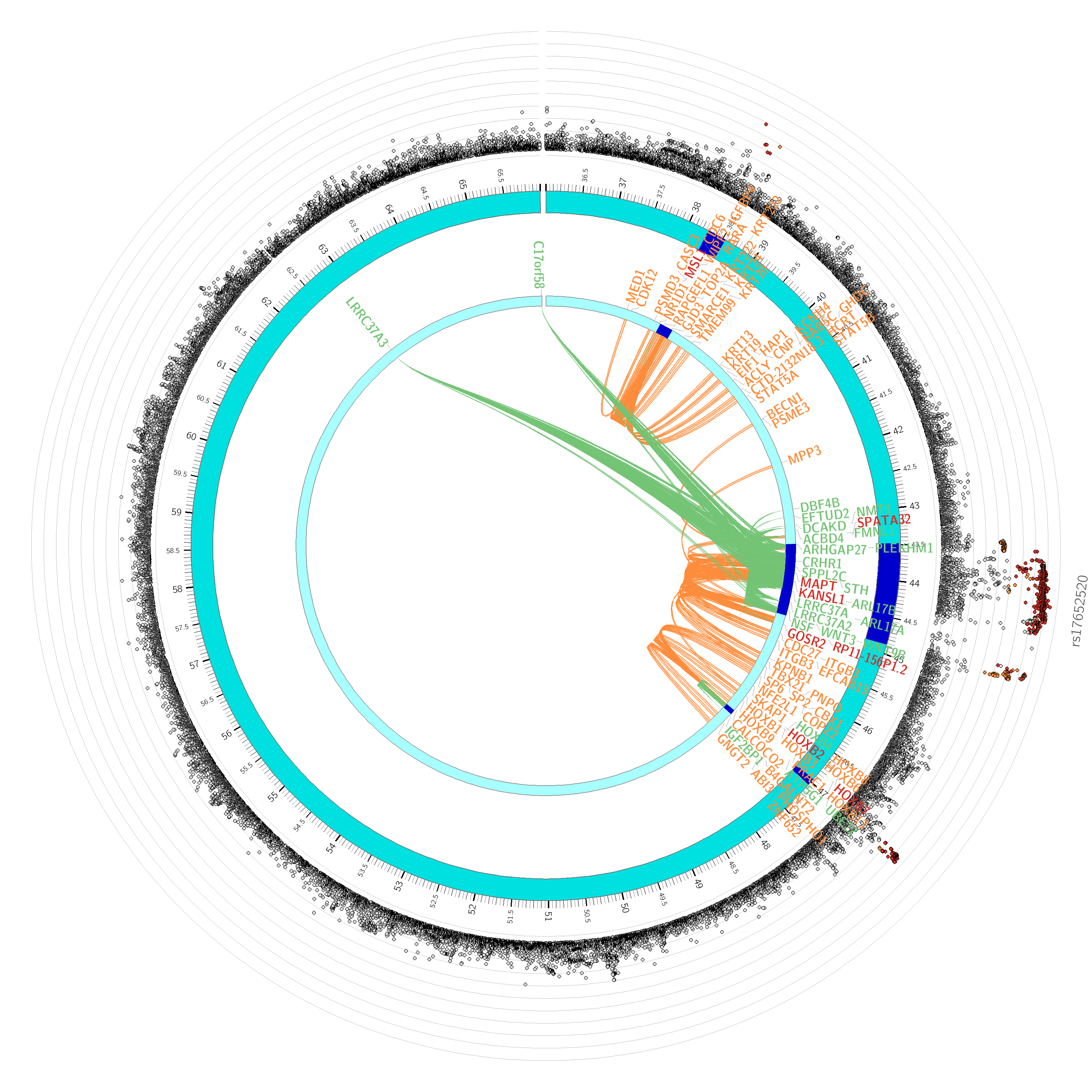

### Supplementary Figure 1K Chromosome 18

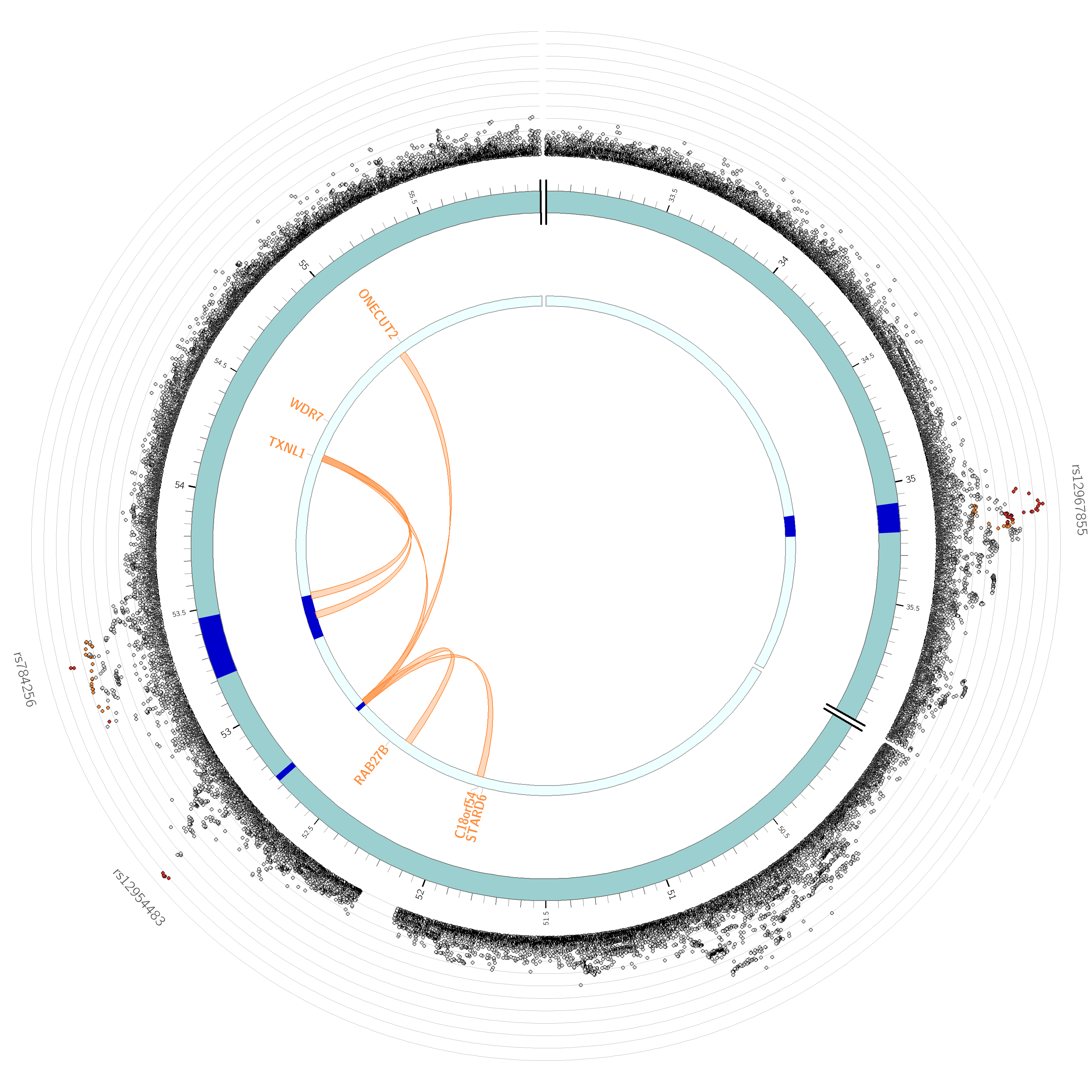

### Supplementary Figure 1L Chromosome 19

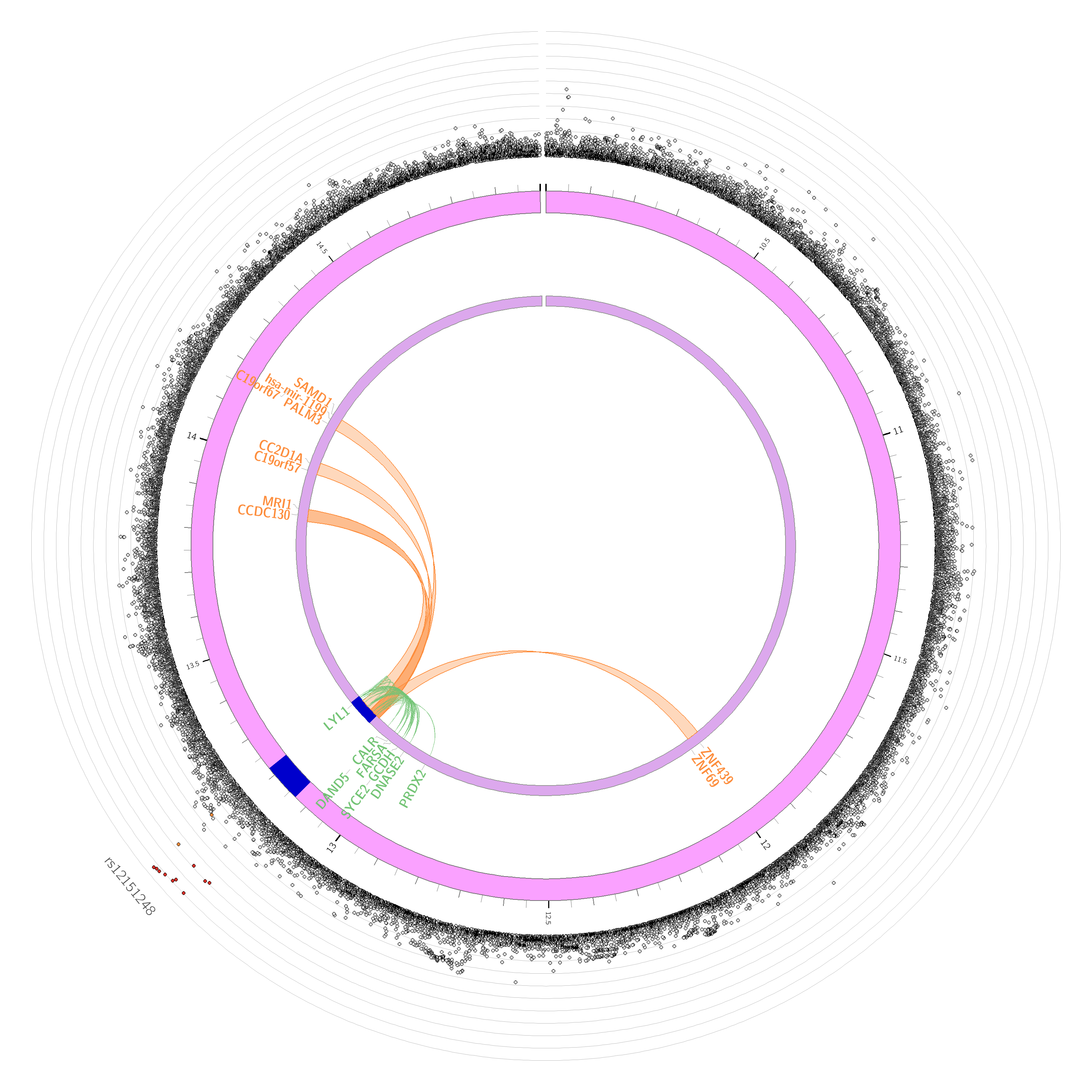

### Supplementary Figure 1M Chromosome 20

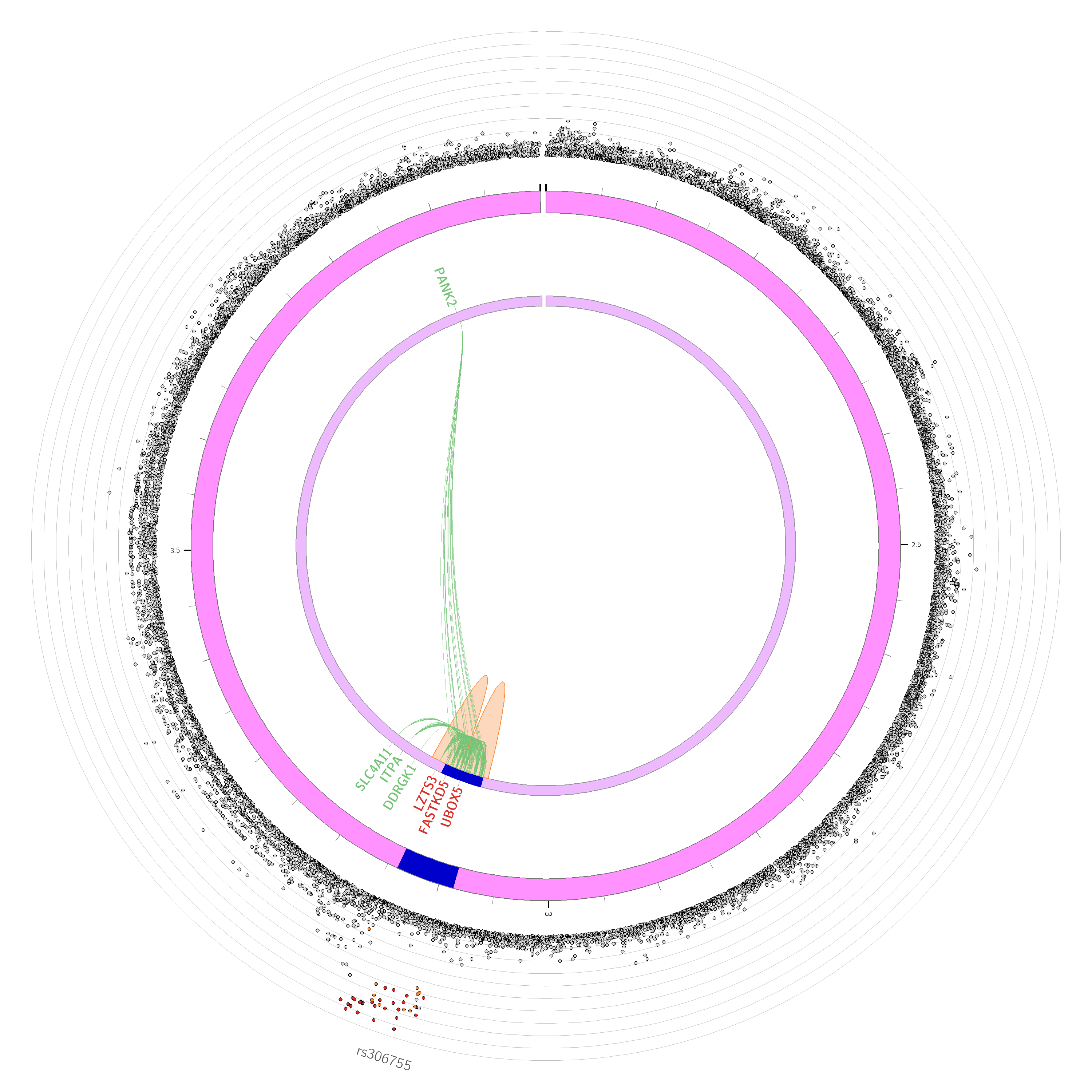
