## Supplementary Figure Captions for "Genetic analysis identifies molecular systems and biological pathways associated with household income"

**Figure legends**

**Supplementary Figure 1.** Circos plots by chromosome illustrating genome-wide significant loci associated with income (Supplementary Figure 1. The most outer layer shows the Manhattan plot and only SNPs where P <0.05 are shown. Each of the SNPs in the genomic risk loci are colour coded indicating the maximum r^2^ with one of the independent significant SNPs in the locus with red indicating the highest r^2^ and blue the lowest r^2^(red r^2^>0.8, orange r^2^>0.6, green r^2^>0.4, and blue r^2^>0.2). SNPs shown in grey are not in LD with any of the genome wide significant SNPs. The rsID of the most significant lead SNP in each loci is shown. The second layer is the chromosomal ring with the independent genomic risk loci highlighted in blue. Next, the genes mapped by chromatin interactions or eQTLs are displayed. Genes mapped using chromatin interactions the gene is displayed in orange, with genes mapped by eQTL shown in green. Genes that are displayed in red are those mapped using both chromatin interactions and eQTLs. Chromatin interaction links (coloured orange for chromatin interactions and green for eQTLs are displayed.

**Supplementary Figure 2A**. Manhattan plot for MTAG derived income phenotype; negative log10 transformed P-values for each SNP are plotted against chromosomal location. The red line indicates genome-wide significance and the black line indicates suggestive associations. **Supplementary Figure 2B**. Functional annotation carried out on the independent genomic loci identified. The percentage of SNPs found in each of the nine functional categories is listed. **Supplementary Figure 2C**. The percentage of SNPs from the independent genomic loci that fell into each of the Regulome DB scores categories. A lower score indicates greater evidence for that SNPs involvement in gene regulation. **Supplementary Figure 2D**. The percentage of SNPs within the independent genomic loci plotted against the minimum chromatic state for 127 tissue/cell types. **Supplementary Figure 2E**. Venn diagram illustrating the overlap of the genes implicated using positional mapping, eQTL mapping, chromatin interaction mapping, that was conducted on the independent significant loci identified in the SNP-based GWAS. Also shown is how these implicated genes overlap with those identified using the gene-based statistics derived using MAGMA.

**Supplementary Figure 3A**. A Manhattan plot of the MTAG derived income phenotype using the gene-based statistics derived using MAGMA; negative log10 transformed P-values for each gene are plotted against chromosomal location. The red line indicates genome-wide significance. **Supplementary Figure 3B.** Enrichment analysis for the MTAG derived income phenotype using the 52 functional categories (27 categories describing enrichment within these categories is shown. The full results including the same functional categories but with a 500kb boundary can be found in **Supplementary Table 26**). The enrichment statistic is the proportion of heritability found in each functional group divided by the proportion of SNPS in each group (Pr(h^2^)/Pr(SNPs)). The dashed line indicates no enrichment found when Pr(h^2^)/Pr(SNPs) = 1. A Bonferroni correction controlling for 52 tests was used to ascertain statistical significance which is indicated by asterisk. **Supplementary Figure 3C.**  Cell type enrichment analysis of the MTAG derived income phenotype. Here, an enrichment of heritability was examined in 10 tissue types. A Bonferroni correction was used to control for the number of independent tests conducted an asterisk indicates statistical significance.
