## Supplementary Results for "Genetic analysis identifies molecular systems and biological pathways associated with household income"

**Functional annotation and gene based analysis of MTAG analysis of income**

Functional annotation of the loci identified using MTAG proceeded using the same methods used to investigate the income phenotype from UK Biobank.^1^ Firstly, it was found that 53.9% of the SNPs in the independent genomic loci were found in intronic regions and the 25.7% were found to be in intergenic regions with 14.6% being ncRNA intronic SNPs. As found in the analysis conducted without MTAG, these independent genomic loci are also predominantly found in regions of the genome involved in gene expression as indicated by 32.13% of the SNPs have an RDB score of <2 and 80.12% have a minimum chromatin state of <8 (**Supplementary Figure 2B-D** & **Supplementary Table 18**).

By examining these loci for overlap with those in GWAS Catalog we found that the loci identified using MTAG overlapped with those previously associated with intelligence (37 loci) and with education (65 loci). In addition we found that the loci linked to income were also associated with coronary artery disease and blood pressure phenotypes (7 loci) as well as with neuroticism (9 loci). Eleven loci previously linked to schizophrenia were linked to the loci associated with income and 2 loci were also linked with subcortical brain region volumes (**Supplementary Table 20**).

Gene-prioritization was also conducted on the income phenotype derived using MTAG. Here, a total of 1,317 genes were identified (**Supplementary Figure 2E** & **Supplementary Table 21**). A total of 392 genes were implicated using positional mapping with 529 implicated using eQTL analysis, with 1,006 genes identified using chromatin interaction mapping. 448 genes were implicated using two mapping strategies with 162 being identified using all three. Of the genes, 162 were identified across all three prioritization methods; 7 were of additional note as they are implicated through chromatin interactions between two independent genomic risk loci. These genes included *ORC3* on chromosome 6, a gene known with links to neuronal proliferation, which was found implicated in a cross locus interaction in the tissues of hESC, IMR90, the Liver, Mesenchymal stem cell, and Mesendoderm tissues. *ZNF589*, *NRBF2*, and *DPYD* were also implicated by all three mapping strategies and chromatin interactions between two independent genomic risk loci in at least five tissue types (**Supplementary Table 22**).

Using a gene-based GWAS conducted in MAGMA, 448 genes attained statistical significance after controlling for multiple tests (**Supplementary Figure 3A** & **Supplementary Table 23**), and 101 genes were implicated by all three mapping strategies and by the MAGMA gene based GWAS.

Two gene sets were significant following correction for multiple testing neurogenesis (gene-set size = 1,337, P = 1.67 × 10^−6^) and reactome pre notch transcription and translation gene set (gene-set size = 25, P = 3.99 × 10^−6^) (**Supplementary Table 24**). The neurogenesis gene-set has been previously linked to intelligence^2^ and neuroticism,^3^ two phenotypes that show genetic correlations with income. This shared association between neuroticism, intelligence, and income, may therefore represent a biological system linked to all three phenotypes. It should be noted that this link between neurogenesis, neuroticism, intelligence, and income may be a case of mediated, or vertical, pleiotropy.^4^

Gene-property analysis conducted on the MTAG-income phenotype replicated the link between gene expression in the brain and association with income (P = 1.05 × 10^−11^). A novel link was also identified between gene expression in the pituitary gland association with income (P = 7.72 × 10^−11^). The link between expression in the testis and income whilst nominally significant (P = 0.016) did not withstand correction for multiple tests using the MTAG-income phenotype (**Supplementary Figure 3B.** & **Supplementary Table 25**). This relationship between gene expression in the brain with differences in income was evident in twelve cortical tissue groupings the most significant of which were the brain cerebellar hemisphere (P = 3.05 × 10^−12^), brain cerebellum (P = 4.36 × 10^−12^), brain frontal cortex (P = 1.03 × 10^−9^), and the brain anterior cingulate cortex BA24 (P = 4.65 × 10^−8^) (**Supplementary Table 26**). A significant relationship was also found for gene expression in the early mid-prenatal developmental period and income (P = 0.002) (**Supplementary Table 27**) but no link was found with any of the age specific expression groupings (**Supplementary Table 28**).

These relationships between income and gene expression across the cortex are also consistent with previous findings pertaining to intelligence,^2^ and so support the notion that intelligence is an intermediary phenotype between molecular genetic inheritance and individual difference in income.

Partitioned heritability conducted using stratified LDSC produced similar, but more precise, findings to the non-MTAG analysis where a significant enrichment for income was found in evolutionarily conserved regions of the genome (P = 1.99 × 10^−17^). In addition, statistically significant enrichment was also found for the histone marks of H3K9ac peaks (P = 1.02 × 10^−6^), and H3K4me1 peaks (P = 2.54 × 10^−6^) as well is in these histone marks more broadly H3K9ac (P = 1.16 × 10^−5^), and H3K4me1 (P = 1.45 × 10^−4^) indicating a role for gene expression differences involved in income differences. Introns were also significantly enriched for income (P = 1.98 × 10^−6^) as were super enhancers (P = 6.81 × 10^−5^). Cell type enrichment analysis also saw that each of the annotations that was significantly enriched in income was also enriched using the MTAG derived income phenotype with the CNS (P = 2.32 × 10^−22^), adrenal/pancreas (P = 5.69 × 10^−7^), cardiovascular (P = 4.30 × 10^−7^), skeletal muscle (P = 3.08 × 10^−8^), and immune/hematopoietic (P = 0.002) annotations replicating. In addition, the cell type annotation of connective bone was significantly enriched for income (**Supplementary Table 29**).
