## Supplementary material for "Genetic analysis identifies molecular systems and biological pathways associated with household income": FAQ

**FAQs regarding:**

**What did you do in this study?**

We performed a genome-wide association study (GWAS) using data from 286,301 individuals aged 39-73 years from across the United Kingdom. These participants provided DNA data as well as information on their health and wellbeing as part of a study called the UK Biobank. In our GWAS, we examined around 18 million points of DNA variation called single nucleotide polymorphisms (SNPs, pronounced ‘snips’). SNPs are the smallest and most common form of genetic variation found (although they are not the only way in which individuals can differ genetically). We looked at these genetic data in relation to household income: a measure of what social scientists often call socio-economic position.

In the participants who had data on income measures and who had donated genetic material for analysis, we tested whether the level of income varied along with their SNPs. First, we examined if people whose household income was higher were more likely to have particular versions (called an allele) of SNPs in their genome. Second, we looked at how much of the income differences between people could be accounted for by differences across the whole set of SNPs that we examined; that is, we used DNA data to examine the ‘heritability’ of income, as indicated by these SNPs. Third, we combined our genetic data with gene expression data, and information on gene locations and their functions. The goal of these analyses was to understand the likely biological effects of any genetic variants that were linked to income. Fourth, because our previous work examining income and genetics in the UK had indicated that DNA might be linked to income through cognitive ability, in the current study we examined the causal role that intelligence might play in individual differences in income in the UK today.

**What did you find?**

Our results suggest that, as we expected, the majority of the reasons that individuals differ in their level of household income was not genetic and likely to be environmental. However, there was some small association between genetic variation and variation in household income.

We identified 30 regions of the genome (termed loci) that were associated with individual differences in household income (**Figure 3**). The SNPs within these loci showed evidence of being involved in gene-expression differences, and we identified that genes expressed in the brain and the synapse were associated with income (**Figure 4** & **Figure 5**). We also found evidence that genes involved in neurogenesis (**Supplementary results**), the process by which new neurons are formed, were associated with household income. Each individual SNP had only a miniscule association with household income, even though it was statistically significant.

We found that variation across all the SNPs in the DNA from this sample accounted for 7.4% of the variation in household income. The rest of the variation is likely due to environmental factors, types of genetic differences that we didn’t measure, and to errors of measurement. Therefore, the large majority of people’s differences in household income are likely to be environmental in origin, according to these results.

We found that people’s genetic differences that were associated with higher household income (i.e. those that accounted for 7.4% of household income variation) overlapped with the genetic differences that were linked to better longevity, health, wellbeing, and intelligence (**Figure 5C** & **Figure 5D**). We found that the genetic effects linked to higher household income overlapped with the genetic differences that were protective against schizophrenia, ADHD, coronary artery disease, and feelings of tiredness and fatigue (**Figure 5C** & **Figure 5D**). We were able to predict up to 2.5% of the differences in income in an independent UK sample using only DNA (**Figure 6C.**).

We used a technique called Mendelian randomisation that is designed to be able to investigate causality using genetic data. We found evidence that being more intelligent is causally related to having a higher income in the UK at present.

**How could genetic differences be associated with something like household income?**

At first this seems confusing: a UK person’s income is a social variable, influenced by factors such as their parents, the part of the country they happen to live in, how the economy is doing, politics and social policies, health, education, and, of course, luck. How can it be associated with genetic differences?

There are many traits that a person has that could contribute to making them more or less likely to earn a higher income. They might be more highly motivated and conscientious at work, for example, or they might be more intelligent, be less susceptible to illness, or simply more interested in the kinds of jobs that lead to higher wages. Some, or all of these, and other psychological and health factors might be associated to some extent by genetic differences, especially if these genetic variants have effects in the brain or in other parts of the body that are important for maintaining one’s health. In our study we identify intelligence as one of the partly heritable factors that might help explain part of the association between genetic differences and differences in household income.

We emphasize that our results imply that by far the largest influence on people’s income is differences in environmental factors. Our findings also indicate that income attainment is a complex process, associated not just with differences in environmental exposures, but also with a non-uniform distribution of things like cognitive capability (intelligence), which is partly heritable, and for which we found some evidence of its being modestly causal to income (**Figure 1**). Our results make a small start at understanding, in the round, why people differ in household income.

**What’s the point of doing this research?**

The prompt for the study was our interest in health inequalities and how to ameliorate them. It is well known that people with a lower income tend to be at greater risk of poorer health than those from more advantaged social backgrounds. Research into the reasons for these health differences has focussed on environmental explanations. However, previous studies have shown that many common health problems are partly due to the genes that people inherit. A small part of the variation in social factors such as income also appears to be modestly associated with differences in their genes. Therefore, we wanted to investigate whether the associations between income and health might be partly accounted for by people’s genetic differences that are associated with both income and health. If politicians or others wish to make effective policies to reduce the health inequalities between people of different social backgrounds, they need to know what brings about those health inequalities.

**Aren’t these effects just population stratification?**

Population stratification describes the existence of a systematic difference in allele frequencies between sub-groups within a larger population. This, one could argue, could occur in instances where individuals selectively mate with those who earn a similar wage. Should this occur, then, over many generations, genetic drift (a random change in allele frequencies) will lead to allele frequency differences between income groups. The result of population stratification in a GWAS study aiming to identify genetic variants linked to income, such as ours, would mean that the link between DNA and income would be spurious, and not tell us anything about why differences in income occur. Therefore, it was important in our study to take steps to minimize, and to test for, and population stratification effects.

We used two methods to limit the effect that population stratification had on our results. First, we ensured that all our participants originated from a similar ancestral background. In this case all our participants were white British individuals. Second, we controlled for the degree to which genetic clustering occurred by including 40 principal components derived from the genotyped DNA-SNP data in our analysis. By doing this we sought to break the association between SNPs and a phenotype that could have been caused by population stratification.

Finally, we tested for population stratification using a method called linkage disequilibrium score regression (LDSC) (Bulik-Sullivan et al., 2015). LDSC is used to test whether the results of a GWAS are due to population stratification or are due to many variants each associated with a small effect (what is termed polygenicity). In our study we used LDSC regression, and the results indicated that over 92% of the signal we identified in our GWAS on income was due to a large number of genetic variants each exerting a small effect (i.e., a polygenic effect) rather than population stratification.

**Have you found “the money gene”?**

We have not found the ‘money gene’ or ‘genes for income’. Income variation is a complex social measure, with many influences. Some potential influences of income—such as some illnesses and some personal traits—are themselves partly heritable. It is possible that some genetic associations with such factors are also picked up in a GWAS study of income. Therefore, our GWAS results should not be interpreted as indicating that there are close causal links between genetic variations and income differences.

Moreover, there were no ‘genes for income’ in the sense that there were no large associations between genetic variants and income differences. Genetic associations with factors such as household income (or intelligence, or some common illnesses) are composed of many thousands of genetic variants, each with a tiny association that adds up.

**Does this mean that an individual’s level of income is determined at birth?**

Our results don’t imply that an individual is predestined to end up earning a certain amount. We found a small association between genetic and income variation in a very large sample. That small association means that, for the most part, people with similar genes will end up with a range of household incomes.

The finding of genetic associations with a trait does not mean that environmental interventions cannot change them. A classic example of this is the disorder of phenylketonuria (PKU). This disorder is genetic in origin, and results in serious medical problems along with intellectual disability. However, by altering PKU patients’ diet from birth, individuals with this condition can lead lives that are not hampered by the disease, or by a reduction in their cognitive abilities. Or, consider eyesight; having poorer eyesight is partly heritable, but these problems can be solved with an environmental intervention: spectacles. These examples show that, even if a trait is highly genetic (and recall that income most certainly is not), it is a possibility that environmental influences can change it.

**Aren’t genetic effects, however small, the sign of a society that is not meritocratic?**

Imagine that we had found no genetic associations with income. What might this mean? It would indicate that the genetic variants associated with, say, better health, higher intelligence, or greater conscientiousness would not be associated with income. Therefore, our finding of a modest genetic association with income could be viewed as an indicator of the “meritocratic” notion that, at least within the population studied here, that an individual’s level of, perhaps, health/ability/personality—as partly influenced by genetic effects—is associated with income to some extent.
